# Topographically driven microclimatic gradients shape patterns of forest structure, diversity, and composition at a forest-grassland transition zone

**DOI:** 10.1101/2022.09.15.508106

**Authors:** Bailey H. McNichol, Ran Wang, Amanda Hefner, Chris Helzer, Sean M. McMahon, Sabrina E. Russo

## Abstract

1. Globally, forests provide important ecosystem services, but anthropogenic change may shift the boundaries of forested biomes, because small-scale environmental changes govern biome transitions. This is especially true in semi-arid forests, where minor topographic and microclimatic changes influence forest functioning and transitions to open biomes such as grasslands. However, we lack quantitative descriptions of topographically driven microclimate variation and how it shapes forest structure, diversity, and composition in these transition zones.
2. Leveraging a 20.2-ha forest inventory plot (Niobrara plot) at a semi-arid forest-grassland transition zone in the Niobrara River valley (Nebraska, USA), we paired data on abundances and distributions of seedlings, saplings, and adults of woody species with topographic and microclimate data to test the hypothesis that if topographic variation causes variation in microclimate that affects forest function, then forest structure, diversity, and composition should vary significantly with topography and microclimate.
3. Microclimatic variation within the Niobrara plot strongly corresponded with topography, creating a sharp water availability and exposure gradient from the river floodplain to the forest-grassland transition zone. The magnitude of microclimate variation corresponded to that of regional macroclimate variation. Mean soil moisture was 10.2% lower along the higher-elevation transition zone than in the canyon bottoms, corresponding to variation across approximately 2.5 degrees of longitude. Mean air temperature increased by 2.2 °C from the canyon bottoms to upper canyon, corresponding to variation across approximately 3 degrees of latitude.
4. Forest structure, diversity, and composition correlated strongly with topographic and microclimatic gradients. More complex forest structure and higher species richness of adults and saplings occurred in moister, less exposed habitats with steeper slopes and lower elevations, whereas seedling stem density and richness were higher in higher-light, moister habitats at lower elevations. Species occupied well-defined topographic niches, promoting high beta diversity along topographic and microclimatic gradients and high species turnover from the floodplain to the transition zone.
5. *Synthesis*: Microclimatic and topographic variation drive patterns of structure, diversity, and composition in the forests at this forest-grassland transition zone. As the macroclimate becomes warmer and drier, topographically mediated microclimatic refuges supporting diverse, structurally complex forested ecosystems may shrink in semi-arid regions.

## 1. Introduction

Forested biomes are critical providers of ecosystem services and regulators of climate from regional to global scales (Anderson-Teixeira et al., 2013; Aznar-Sánchez et al., 2018; Bonan, 2008; FAO & UNEP, 2020; Hisano et al., 2018; Lawrence & Vandecar, 2015). Forested biomes are commonly thought of as large expanses of forest, but, globally, many forests abut other biomes at transition zones (also known as ecotones), such as the forest-savanna transition zone (Cardoso et al., 2021; Hoffmann et al., 2012a; Oliveras & Malhi, 2016; Veenendaal et al., 2015), the forest-grassland transition zone (Abiem et al., 2020; Barros et al., 2018; Müller et al. 2012), and the forest-marsh transition zone (Hall et al. 2022; Jobe & Gedan, 2021). Forests at biome transition zones combine species from different ecological zones, constituting unique communities (Cardoso et al., 2021; Charles-Dominique et al., 2018; Oliveras & Malhi, 2016). Biome transition zones occur because sufficiently large changes in environmental conditions, including microclimate, fire regimes, and soils, cause shifts in the vegetation type best suited to the local conditions (Allen & Breshears, 1998; Higgins et al., 2016; Pellegrini et al., 2021; Scheiter et al., 2020).

Global models of the earth’s climate have established that for most regions, climate is changing (Masson-Delmotte et al., 2019), but we have limited information on how large-scale climatic shifts will translate into variation in the climatic conditions that trees experience at local scales (hereafter, ‘microclimate’, or climate at the individual to sub-stand scale) (De Frenne & Verheyen, 2016; Hofmeister et al., 2019; Jucker et al., 2020; Pool et al., 1918). Understanding the responses of individual trees and populations to local conditions is crucial, as these responses in aggregate drive forest change (Blonder et al., 2018; Carnicer et al., 2021; Davis et al., 2019; De Frenne et al., 2013; Zellweger et al., 2020). Forests at transition zones may be more susceptible to direct effects of changes in microclimate, as well as its downstream consequences for fire regimes and resilience to pests (Cardoso et al., 2021; Hoffmann et al., 2012a,b; King et al., 2013; Pellegrini et al., 2021), because these are the processes that define the transition between biomes.

In water-limited, semi-arid forests where conditions are challenging for tree growth (Allen & Breshears, 1998; Peters, 2002), small changes in temperature, water availability, and light can affect forest structure (Adams et al., 2014; Andrews et al., 2020; Sun et al., 2019; Szejner et al., 2020). Plant species diversity and composition also vary sharply with resource availability (*e.g.*, light, water, nutrients) along topographic gradients (Adams et al., 2014; Baldeck et al., 2013; Jucker et al., 2018; Laliberté et al., 2014; Liautaud et al., 2020; Russo et al., 2005, 2012). Since more diverse forests provide a broader suite of ecosystem services and have increased resilience and resistance to climate extremes (Isbell et al., 2015; Pardos et al., 2021; Reu et al., 2022; Sakschewski et al., 2016; Tilman et al., 2014), it is essential to understand how diversity is structured by fine-scale environmental conditions. However, many studies use topography as a proxy for microclimatic variation without explicitly quantifying the underlying variation in environmental conditions that affect tree physiology (Méndez-Toribio et al., 2016; Muscarella et al., 2020; Yeakley et al., 1998). In semi-arid forests, this is a critical gap, because water availability defines forest structure and composition. At large scales, macroclimate largely governs biome transitions (Anderson-Teixeira et al., 2015; Brice et al., 2020; Gosz & Sharpe, 1989), but topography can create climatic refugia, altering these transitions at smaller scales (Adams et al., 2014; Dobrowski, 2011). Understanding how topography modulates macroclimate to create the microclimates that plants experience is essential for understanding transition zones between biomes, plant community structure at transition zones, and how both are responding to anthropogenic change.

In this study, we evaluated the extent to which topographically driven variation in microclimate shapes the structure, diversity, and composition of a boundary forest at the forest-grassland transition zone in the Niobrara River valley, Nebraska, USA (Figure 1). Our study site is located at the edge of the Nebraska Sandhills, which is the largest sand dune formation in the Western Hemisphere (Loope & Swinehart, 2000; Sridhar et al., 2006; Tolstead, 1942a) and overlies the High Plains aquifer, one of the most important sources of groundwater in North America (Ajaz et al., 2020). The warm, semi-arid macroclimate constrains the predominant biome of this region to grassland, but forests occur in cooler, moister canyons draining to the Niobrara River (Kantak, 1995; Kaul et al., 1988). Due to the region’s topographic, hydrologic, and hydrogeologic complexity (Hearty, 1978), tree species with predominantly western, boreal, and eastern distributions co-occur (Bessey, 1905; Kaul et al., 1988). The populations of most tree species in the Niobrara forests are therefore not only at the edges of their habitat range in the forest-grassland transition zone, but also at edges of their species’ geographic range (Bessey, 1905; Kantak, 1995; Kaul et al., 1988; Tolstead, 1942b).

**Figure 1.**
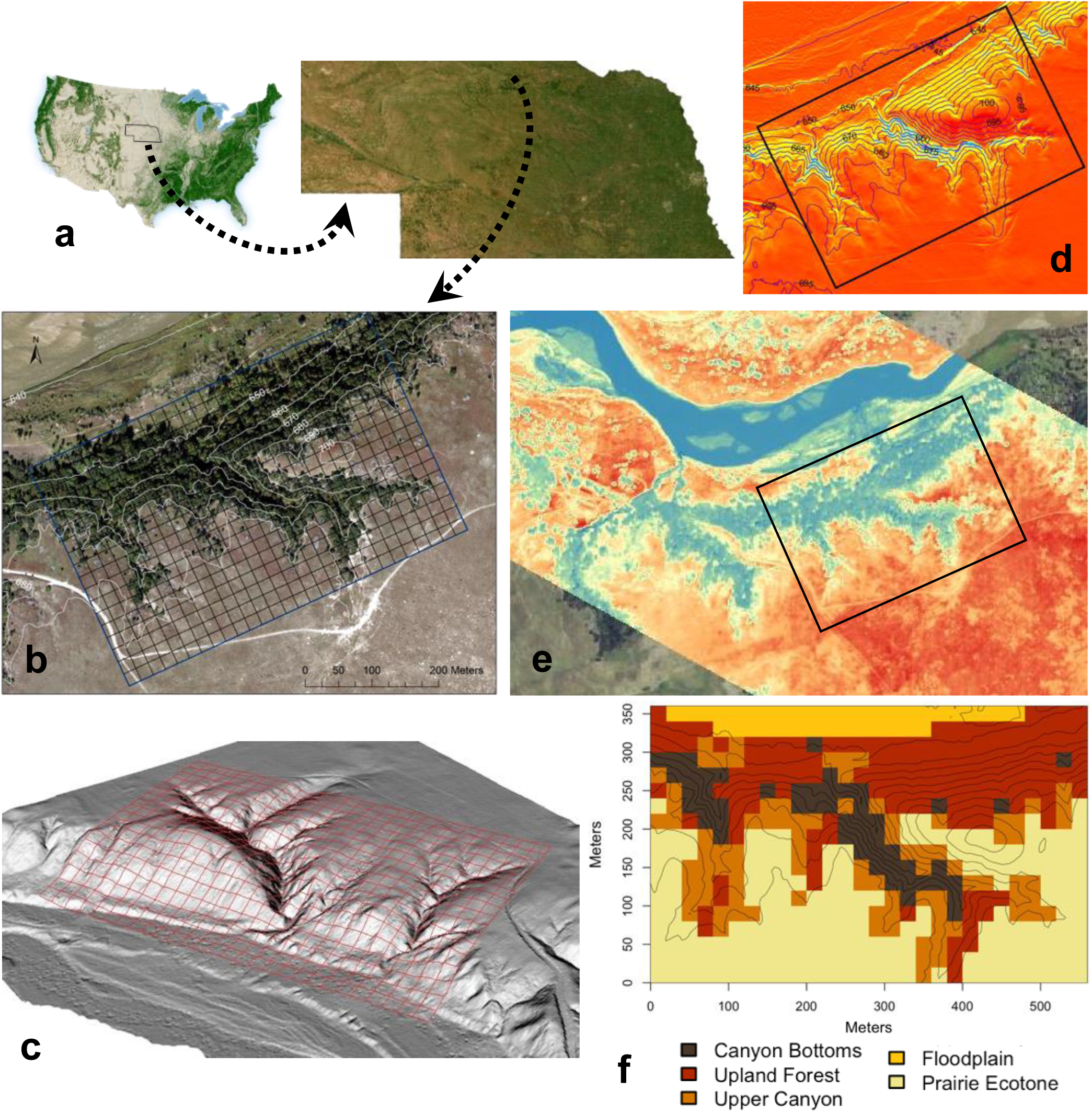
Location, topographic gradients, and habitats of the 20.2 ha Niobrara forest dynamics plot along the forest-grassland biome transition zone in the Niobrara River Valley of Nebraska in the Great Plains, USA. (a) Location of the Niobrara plot, with respect to a map of forest cover in the continental United States (taken from the U.S. Forest Service’s online Forest Atlas of the United States) and a 15-m resolution aerial map of Nebraska (Image source: ESRI, 2022); (b) a high-resolution RGBN image obtained on 18 September 2019, with an UltraCam Eagle camera (Vexcel Imaging), showing the boundary of the Niobrara plot and 20 x 20 m quadrats with 10-m elevation contours; (c) digital elevation model (DEM) used to estimate the five topographic variables (elevation, solar radiation, slope, eastness, and northness), with the Niobrara plot and quadrat boundaries overlaid (U.S. Geological Survey 2017); (d) a map of cumulative annual solar radiation [Wh/m^2^; red-yellow-blue color ramp indicates high to low radiation showing gradients in exposure within the Niobrara plot with respect to elevation contours, based on the DEM shown in (c)]; (e) a thermal image (blue colors depict cooler, and red colors depict hotter, relative temperatures) of the region with the Niobrara plot boundary in black (image obtained on 13 July 2022); (f) a map of habitats in the Niobrara plot, overlaid with 5-m elevation contour lines.

Biome transition zones, such as those in the Niobrara region, are considered indicators of the net effects of anthropogenic change, offering the ability to forewarn how species in the core parts of forested biomes may respond to direct and indirect effects of climate change. However, quantifying the effects of anthropogenic change on these zones is impossible without first understanding the fundamental ecological drivers causing these transitions (Oliveras & Malhi, 2016). In a 20.2-ha forest inventory plot along the Niobrara forest-grassland transition zone sampling seedlings to adults for a total of 37 woody species, we coupled data on woody species’ abundances and spatial distributions, topographic data on elevation, slope, aspect, and cumulative annual solar radiation derived from a digital elevation model, and locally sampled microclimate data (understory photosynthetic photon flux density, air temperature and relative humidity, vapor pressure deficit, and soil temperature and moisture) to identify the predominant abiotic drivers of variation in forest structure, diversity, and composition. We tested the hypothesis that if variation in topography causes spatial and temporal variation in microclimate affecting forest functioning across the transition zone, then there should be strong relationships of both topography and microclimate with forest structure, diversity, and composition across tree size classes.

## 2. Methods and Materials

### 2.1 Study site and census methods

The Niobrara Forest Dynamics Plot (hereafter, Niobrara plot; 42°46’48.83” N, 100°01’15.56” W; Figure 1) is part of the Smithsonian Forest Global Earth Observatory (ForestGEO) network of forest dynamics plots (https://forestgeo.si.edu/sites/north-america/niobrara), which quantify forest diversity and dynamics worldwide. The 20.2 ha (560 m x 360 m) Niobrara plot was established in 2019 using standard ForestGEO methods (Condit, 1998) within the Niobrara Valley Preserve (Johnstown, Nebraska, USA) in the North American Great Plains (Figure 1a, b). The Niobrara Valley Preserve is owned and managed by The Nature Conservancy. They manage for expansion of *Juniperus virginiana*, which is occurring throughout the Great Plains (Meneguzzo & Liknes, 2015; Msanne et al., 2017; Twidwell et al., 2013), paralleling woody encroachment globally (Abiem et al., 2020; Cardoso et al., 2021; Pálinkás, 2018). Management activities in the Niobrara plot consisted of a thinning of large *J. virginiana* stems in summer 2017 and a low-intensity prescribed burn in October 2020.

From 2000-2015, the average cumulative annual precipitation on the Niobrara plot was 556 mm, most of which occurred during the warmest part (May-August) of the growing season (April-October) between. The mean annual temperature was 9.6 °C, and average minimum and maximum daily temperatures were −11.3 and 32 °C, respectively (PRISM Climate Group, 2020). This climate regime lies within the temperate grassland biome, near the boundary with the woodland/shrubland biome (Chapin et al. 2011; Whittaker, 1975). Soils in the plot are entisols, which are characterized by little development of soil profile horizons and are commonly found at sites of recently deposited materials, or of parent materials resistant to weathering (*e.g.*, sand) (Soil Survey Staff, USDA NRCS, 2015). Inavale and Almeria loamy fine sands occur along the floodplain (entisols, 0-2% slopes), McKelvie-Fishberry-Rock outcrop complex occur in the forest and along the ecotone (loamy entisols typical on valley sides, 11-60% slopes), and the Valentine-Simeon complex (excessively drained sandy entisols, 9-40% slopes) are found in the grassland (Soil Survey Staff, USDA NRCS, 2019).

The forests along the south side of the Niobrara River valley, which the Niobrara plot samples, have a more sheltered north-facing aspect. To the south, they are bordered by Sandhills prairie, creating a dramatic forest-grassland transition zone (Kaul et al., 1988; Figure 1b) with well-defined thermal environments visible at large spatial scales (Figure 1e). The Niobrara plot encompasses a 59-m elevational gradient (644-703 m.a.s.l.) from the Niobrara River floodplain to the transition zone at higher elevations (Figure 1b, c). Forests occur in and along the canyons and slopes draining into the Niobrara River and are supported by groundwater seeps and springs that emerge along the canyon walls (Hearty 1978), contributing to a cooler, moister microclimate (Kaul et al., 1988; Tolstead, 1942a; Figure 1c, d, e). As a result of these local microclimatic conditions, many plant species’ geographic ranges extend along the Niobrara valley, causing these species to be at range edges. The forests consist of an unusual combination of westerly (*Pinus ponderosa*), easterly (*e.g.*, *Juglans nigra*, *Tilia americana*), boreal (*Betula papyrifera*), and centrally distributed species (*e.g.*, *Quercus macrocarpa*), and widespread species (*e.g.*, *Rhus glabra*) (Table S1).

The Niobrara plot is subdivided into a grid of 504 20 × 20 m quadrats (hereafter, quadrats; Figure 1b, c) permanently marked by posts georeferenced to within *ca.* 10 cm by a professional surveyor. In the first plot census (2018-2019), within each quadrat, all woody stems (trees, shrubs, and lianas) ≥1 cm in stem diameter at breast height (1.3 m, DBH) were individually tagged, identified to species, measured for DBH, mapped using a total station (Leica Flexline TS06 Plus, Leica Geosystems AG), and georeferenced based on surveyed posts. Secondary stems (*i.e.*, multiple stems on the same individual emerging from the main stem below 1.3 m and with DBH ≥1 cm) were also tagged for better estimation of basal area and aboveground biomass (AGB). In the first census, 339 of the 504 quadrats in the plot (67%) had at least one woody stem ≥ 1 cm, for a total of 8,299 stems of 27 woody species, including 25 deciduous broadleaved species (three lianas, four shrubs, and 18 trees) and two evergreen coniferous tree species (*J. virginiana* and *P. ponderosa*) (Table S1).

In May-July 2021, 1 m x 1 m permanently marked seedling subplots (hereafter, subplots) were established at the northeastern-most corner post of each of the 504 20 x 20 m quadrats in the Niobrara plot (sampling 0.0504 ha of the Niobrara plot). All seedlings of woody species (stem < 1 cm in DBH) in each subplot were tagged and identified to species. During the first seedling census, 290 subplots of the 504 subplots (58%) had at least one woody seedling, for a total of 2,196 seedlings of 31 woody species, including 29 deciduous broadleaved species (five lianas, nine shrubs, and 15 trees) and two evergreen conifers (*J. virginiana* and *P. ponderosa*) (Table S1). Thus, there were a combined total of 37 woody species recorded in the first tree and seedling censuses in the Niobrara plot. The liana *Parthenocissus quinquefolia* was excluded from analyses due to difficulty in quantifying the number of individuals.

### 2.2 Estimation of DEM-derived topographic variables

We obtained a 1-m resolution Digital Elevation Model (DEM) derived from remotely sensed Light Detection and Ranging (LiDAR) data for Brown County, Nebraska (U.S. Geological Survey, 2017; Figure 1c). We used the DEM encompassing the Niobrara plot to estimate the following five topographic variables: elevation (m), slope (%), aspect (degrees), and the cumulative annual solar radiation (Wh/m^2^; Figure 1b-d) (following Fu & Rich, 2002), using ESRI ArcMap version 10.8 (ESRI, 2020). Aspect, a circular variable that varies from 0-360 degrees, was decomposed into two linear variables representing “northness” and “eastness” by converting to radians, and taking the cosine to derive northness (where −1 is due south, 1 is due north, and 0 represents either east or west), and the sin to derive eastness (where −1 is due west, 1 is due east, and 0 represents north or south) (Gillingham & Parker, 2008; Roberts, 1986). Mean values of each topographic variable were obtained for each quadrat by averaging the 1-m scale values from the DEM encompassed within a quadrat. The 20-m scale is appropriate for linking variation in topography with forest structure, diversity, and composition because larger scales would encompass too much topographic variation, but smaller scales would not encompass enough woody stems for robust analyses. Values for each topographic variable for every woody stem in the plot were estimated by matching the coordinates of the stem to the nearest coordinate from the 1-m scale DEM (ESRI, 2020).

### 2.3 Monitoring of microclimate conditions along environmental gradients

We quantified variation in microclimate conditions throughout the 2021 growing season using ten monitoring stations deployed along the topographic gradients in the Niobrara plot from April to November 2021 (full leaf-out occurred in late May; Stations 1-10 in Figure S1). At each station, understory light availability (photosynthetic photon flux density; PPFD, *μ*mol/m^2^/s;) was measured with 1-2 quantum sensors (Li-Cor LI-190R) and air temperature (°C) and relative humidity (RH, %) were measured at approximately 1 m height above the ground (Campbell Scientific HMP35C, CS-215, and HygroVUE5). Soil temperature (°C) was measured at a depth of approximately 10 cm (Campbell Scientific 105E-L and 108), and surface soil moisture (volumetric water content; VWC, %) was measured with 1-2 time-domain reflectometry sensors to a depth of 30 cm (Campbell Scientific CS616). Additionally, at the station in grassland, just past the forest-grassland transition (hereafter referred to as the prairie ecotone; Station 10), an anemometer (Campbell Scientific 014A-L) and tipping bucket (Texas Electronics model TR-525I) were deployed to measure wind speed (m/s) and rainfall (cm). Mean values over fine timescales (every 1 min for light and every 5 min for all other variables) were estimated based on temporarily stored measurements every 5 s and recorded on battery-powered data loggers (Campbell Scientific models CR-800 and CR-1000).

The microclimate stations measured conditions at fine time scales, but at only ten locations in the Niobrara plot (Figure S1). To quantify variation in surface soil moisture with greater spatial coverage along the topographic gradients, manual point measurements of VWC (%) were taken using a hand-held time-domain reflectometer (Campbell Scientific Hydrosense II) in 73 quadrats distributed across each habitat (canyon bottoms = 15, upland forest = 14, upper canyon = 14, floodplain = 15, prairie ecotone = 15; see *Methods* 2.4 and Appendix S1 for habitat definitions). Measurements at approximately the same 3 locations per quadrat were taken on a rain-free day six times between May and October 2021, and quadrat sampling order was varied to avoid confounding time of day with habitat-related variation.

### 2.4 Statistical analysis

Our hypothesis leads to the following four predictions, which we evaluated using statistical analyses. (P1) Forest habitats can be derived based on topographic variation, and there should be strong relationships of both habitat-based categorical and continuous topographic variables with spatial and temporal variation in microclimate (means and/or coefficients of variation). (P2) Topographic variation should correlate with forest structure, diversity, and composition for seedlings and trees of woody species, using both habitat-based categorical analyses and continuous analyses with the original topographic variables. (P3) The correspondence between topography and forest structure, diversity, and composition should be reflected in distinct topographic niches of woody species. (P4) Forest composition should be correlated with microclimate variation. All analyses were conducted in R software version 4.0.2 (R Core Team, 2020).

#### Derivation of habitat types

Following previous studies (*e.g.*, Kenfack et al., 2014; Valencia et al., 2004), we defined five categorical topographic habitats using the five topographic variables (detailed description in Appendix S1). To capture the covariation among these variables, we conducted principal components analysis (PCA), and found the first principal component (PC1; 42.7% of variation explained) was correlated with variation in slope, solar radiation, and elevation, and PC2 (22.6% of variation explained) was correlated with aspect (Figure S2a, 1f; Table S2). Habitats were defined based on cutoffs for the quadrats’ scores for PC1 and PC2. The distribution of quadrats assigned to each habitat across the Niobrara plot is shown in Figure 1f (canyon bottoms: 2.04 ha; upland forest: 5.56 ha; upper canyon: 3.56 ha; floodplain: 1.48 ha; prairie ecotone: 7.52 ha).

#### P1 Quantifying microclimate variation across habitats and along the topographic gradient

We derived the vapor pressure deficit (VPD, kPa) for each measurement day from the air temperature and relative humidity, using air temperature to calculate the dewpoint temperature, and then calculating VPD using Tetens’ formula as *e_s_* − *e_a_*, where *e_s_* is the vapor pressure at the current air temperature and *e_a_* is the vapor pressure at dewpoint temperature (Monteith and Unsworth, 2008). Daily microclimate data collected at the ten stations (Table S3) were summarized across all measurement days based on means and standard deviations (SD) using the ‘chron’ (James & Hornik, 2020), ‘dplyr’ (Wickham et al., 2021), ‘lubridate’ (Grolemund & Wickham, 2011) and ‘padr’ packages (Thoen, 2021). To examine variation in microclimate between stations, between habitat types, and across the growing season, we calculated the monthly mean values and coefficients of variation (SD/mean*100; CV) for each variable at each station. For missing monthly values at eight stations in April and May for RH, air temp, and VPD due to delayed sensor deployment, we imputed values using the *imputePCA* function (‘missMDA’ package; Josse & Husson, 2016). We ran separate PCAs on the scaled monthly microclimate means and CVs for each station, as well as corresponding permutational multivariate analyses of variance (perMANOVA; Anderson et al., 2006) assessing the effect of habitat on the means and CVs using the Gower distance metric (Gower, 1971), with month nested within habitat and a block effect to account for variation among stations within the same habitat. perMANOVA was implemented using *adonis2* (‘vegan’ package; Oksanen et al. 2020). To assess the degree to which microclimatic variation is driven by topographic variation, we extracted the first three PCs from the PCA on topographic variables that was used to define categorical habitat types (Appendix S1) and conducted perMANOVA to assess the additive effects of topographic PC1, PC2, PC3, and month on the dissimilarity (Gower) between stations in each month in their microclimate means and CVs.

To assess variation in surface soil VWC between habitat types and across the growing season, we fit a linear mixed effects model on the manually measured soil VWC data, with habitat and sampling date as fixed effects and quadrat as a random effect, using the *lme* and *anova.lme* functions (‘nlme’ package; Pinheiro et al., 2021). To assess goodness of fit, we calculated pseudo-*R^2^* values (*pR*^2^) to quantify variance explained by the fixed effects alone (marginal *pR*^2^) and by both the fixed and random effects (conditional *pR*^2^) using the *r.squaredGLMM* function (‘MuMIn’ package; Bartón, 2022). To determine whether there were differences in VWC between habitats and dates, we used post-hoc comparisons of means using the *glht* function with a single-step adjustment for multiple comparisons (‘multcomp’ package; Hothorn et al., 2008).

To explore topographic constraints on variation in soil moisture, we fit quantile regressions (Cade & Guo, 2000) with the manually measured VWC as the response variable and the additive effects of sampling date and one of either elevation, slope, or solar radiation as predictors in three separate models. Quantile regression weights observations depending on the quantile of the response variable and is useful when data show boundary-type relationships (Koenker & Bassett, 1978; Koenker & Hallock, 2001). The quantile regressions were implemented using the *rq* function with *τ* = 0.5 (median quantile) and *τ* = 0.95 (95^th^ quantile) using the Barrodale and Roberts algorithm (‘quantreg’ package; Koenker, 2021). To assess the goodness of fit of these models, we calculated *pR*^2^ values following Koenker & Machado (1999).

#### P2 Modeling topography-driven variation in forest structure, diversity, and composition

To assess patterns of forest structure and diversity along the topographic gradients, we ran separate analyses on (a) adults and saplings (referred to together hereafter as ‘trees’; DBH ≥1 cm), and (b) seedlings (DBH < 1 cm) of woody species. To quantify forest structure and diversity, we calculated tree stem density, total basal area, total AGB, and species richness and diversity for trees in each quadrat using functions in the ‘fgeo’ (Lepore et al. 2019), ‘allodb’ (Gonzalez-Akre et al., 2021), and ‘vegan’ (Oksanen et al. 2020) packages. Stem density was the number of individual trees per quadrat. Several species often form individuals with multiple large stems (*e.g.*, *T. americana*), so calculations of total basal area and AGB included multiple stems. Total quadrat basal area (m^2^) was calculated from DBH (cm) as Σ *π* × (DBH)^2^/(4×10,000 cm/m^2^). The ‘allodb’ package uses a database of species and genus-specific allometric equations and wood densities, combined with DBH measurements, to estimate the AGB of all stems of an individual in kg (Gonzalez-Akre et al., 2021), which were summed for trees in each quadrat to obtain the quadrat-level AGB. We scaled tree stem density, basal area, and AGB to a per ha basis. Finally, we calculated woody species richness (number of species per quadrat, or per 400 m^2^) and Shannon’s diversity as *H* = −Σ *p_i_* × ln (*p_i_*), where *p_i_* is the proportion of individuals in the *i*th species (Magurran, 2004). Analyses of trees included quadrats with at least one woody stem ≥1 cm in DBH in the first census (*N*=339 of 504 quadrats). To assess structure and diversity in the seedling size class, within each seedling subplot, we calculated stem density (number of woody seedlings), species richness (number of woody species), and Shannon’s diversity. Analyses of seedlings included all subplots with at least one woody seedling in the first census (*N*=290 of 504 subplots).

We used two modeling approaches to quantify topography-driven variation in forest structure and diversity in the tree and seedling size classes in separate models. First, we used linear models with habitat (canyon bottoms, upland forest, upper canyon, floodplain, prairie ecotone) as a categorical fixed-effect predictor fit using ordinary least squares regression and quadrat as the unit of replication. Box Cox transformations were applied to response variables to meet assumptions of homoscedasticity and normality (Venables and Ripley, 2002). When there was a significant omnibus test for habitat, pairwise differences between habitats were evaluated via post-hoc multiple comparisons using Tukey’s Honestly Significant Differences (HSD) with a 5% family-wise error rate, using the *TukeyHSD* function.

Second, we used generalized additive models (GAM) with the original topographic variables as continuous predictors to assess their main and interactive effects on tree (or seedling) structure and diversity and with quadrat (or subplot) as the unit of replication. GAMs are a non-parametric extension of linear regression that apply an additive smoothing function to the predictor variables and take the following form: *g*(*E*(*Y*)) = *b* + *s*_1_(*X*_1_) + … + *s_p_*(*X_p_*), where *E(Y)* is the response variable, *g(Y)* is the link function, *b* is the intercept, *s(X)* is the smoothing function of each of the predictor variables, and *X_p_* are the *p* continuous predictor variables (Hastie & Tibshirani, 1987). We used GAMs because they flexibly accommodate nonlinear relationships, while also avoiding model overfitting by penalizing the smoothing parameter (Yee & Mitchell, 1991). To account for mild spatial autocorrelation identified in exploratory analysis (assessed using *augment* in the ‘broom’ package; Robinson et al., 2021), we included a two-dimensional thin plate spline of the quadrat coordinates in each GAM (Hutchinson & Gessler, 1994). We fit GAMs with a Gaussian error distribution and identity link function using the *gam* function (‘mgcv’ package; Wood, 2011). Elevation was standardized by subtracting the minimum elevation from each value prior to analyses.

We used model selection to find the most-supported model among 59 candidate GAMs fitted for each response variable (Table S4). None of the three-way interactions between the five topographic variables were significant, so the most complex model (global model) included all two-way interactions between the five topographic variables, and the simplest (null model) was the intercept only model. There was no evidence of multicollinearity between topographic variables, as all variance inflation factors in the global model were < 2.5. Box Cox transformations were applied to meet the assumptions of homoscedasticity and normality. We performed model selection using Akaike’s Information Criterion (AIC) and determined the relative likelihood of candidate models using Akaike weights (Akaike, 1973; Burnham et al., 2002), using the *map* and *map_dbl* functions (‘purrr’ package; Henry and Wickham, 2020) and the *pander* function (‘pander’ package; Daróczi and Tsegelskyi, 2021) to compare models. We plotted model predictions for the most-supported model, setting predictors not plotted on the x-axis at either their first and third quartile values (elevation) or their minimum value (northness and eastness).

We quantified variation in species composition for trees with respect to both categorical habitat types and continuous topographic variables. We fit a perMANOVA using Cao dissimilarity (Cao et al., 1997) between quadrat pairs as the response variable and habitat type as the predictor. To test whether there was greater compositional overlap within than between habitats, we used analysis of similarity on quadrat-level Cao dissimilarity using the *anosim* function in ‘vegan’. To test whether habitats differed in the variance in species composition, we calculated the within-habitat multivariate dispersion for each habitat using the *betadisper* function in ‘vegan’ and compared it between habitats using the *anova* function with post-hoc multiple comparisons (Tukey’s HSD).

To assess variation in community composition between quadrats along the topographic gradients, we used non-metric multidimensional scaling (NMDS) analyses on the between-quadrat dissimilarities (Cao) with three dimensions (k = 3) and a maximum of 999 iterations, using the *metaMDS* function in ‘vegan’. To test the influence of geographic distance on species composition, we conducted multiple regression with matrices (MRM) using the *MRM* function (‘ecodist’ package; Goslee & Urban, 2007) with Cao dissimilarity in composition as the response variable and the matrix of geographic distances between each quadrat pair as the predictor. Because geographic distance only explained 2.0% of the variation, we excluded this variable from further analyses of composition. To assess the effects of topography on composition, we conducted MRM with the between-quadrat Gower dissimilarities of all topographic variables as predictors and the Cao dissimilarities in composition as the response variable. Topographic variables were scaled by dividing by the standard deviation. To partition variance explained by each topographic variable, we estimated adjusted *R*^2^ from a linear model with the Gower dissimilarities in all topographic variables as predictors (full model) and linear models excluding each one of the variables, and then subtracted their adjusted *R*^2^ values from that of the full model (Swenson, 2014).

#### P3 Topographic niches of woody species

To assess the degree to which woody species (with >1 individual in the Niobrara plot) occupy distinct topographic niches, we determined the elevation (m) and cumulative annual solar radiation (Wh/m^2^) for each woody stem ≥1 cm in DBH, and constructed species-level boxplots for both variables, ordering species by their median values. We assessed the density of individuals and outliers beyond the first and third quartiles ± 1.5 times the interquartile range for each species.

#### P4 Assessing the influence of microclimatic variation on forest composition

We directly analyzed the relationship between variation in microclimate and species composition using perMANOVA and MRM. To quantify species composition of trees near each of the nine forested microclimate stations (excluding Station 10 in the grassland), we defined a 10-m radius around each station, ensuring the entire area was within the habitat type sampled by that station, and estimated Cao dissimilarities between pairs of areas. We calculated station-level microclimate means and CVs (one value corresponding to each station, as required for analyses) by averaging across monthly values for VPD and soil temperature and moisture (scaled by dividing by the SD). Due to the limited number of stations, we elected to include VPD, but not air temperature nor RH, in analyses, as VPD is derived from and hence captures variation in these two variables, as evidenced by the strong correlation of VPD with RH (*r* = −0.99) and air temperature (*r* = 0.88). Additionally, due to the potential for multicollinearity between VPD and PPFD (*r* = −0.47), and the fact that measurements of PPFD only represent understory light availability, we excluded PPFD from analyses. We used separate perMANOVAs to assess the effects of the microclimate means and CVs on species composition. We also calculated Gower dissimilarities between stations using the microclimate means and CVs and tested their relationship with the Cao dissimilarity in composition using MRM.

## 3. Results

### 3.1 P1 Strong microclimatic variation between habitats and along the topographic gradient

At the prairie ecotone, daily rainfall averaged 0.17 ± 0.46 cm (SD), mostly occurring in large rainstorm events throughout the growing season, and the cumulative rainfall between 21^st^ of May to the 6^th^ of November 2021 was 28.1 cm (Figure S3a). Windspeeds averaged 2.6 ± 1.2 m/s, with maximum windspeeds up to 17.4 m/s (Figure S3b). Microclimate conditions monitored at all stations varied strongly between habitats, and especially across the forest-grassland transition zone (Figure 2; Table S3), which is clearly visible in the thermal remote sensing image (Figure 1e). Light availability (PPFD) was considerably higher in the floodplain and prairie ecotone compared to other habitats in all months (Figure 2a). Air temperature was the warmest in the prairie ecotone from June onwards and was considerably cooler in the canyon bottoms compared to other habitats throughout the growing season (Figure 2b). RH was highest in the canyon bottoms between April-July, and in the floodplain from August-November, whereas the prairie ecotone experienced the lowest RH in all months (Figure 2c). VPD was highest in the prairie ecotone throughout the growing season, and consistently lowest in the moist canyon bottoms and floodplain (Figure 2d). As with air temp, surface soil temperature was consistently highest along the exposed prairie ecotone and lowest in the canyon bottoms from May-September (Figure 2e). Surface soil VWC was highest throughout the growing season in the upper canyon (Station 5), which was situated close to a spring, followed by the canyon bottoms and then the floodplain (Figure 2f).

**Figure 2.**
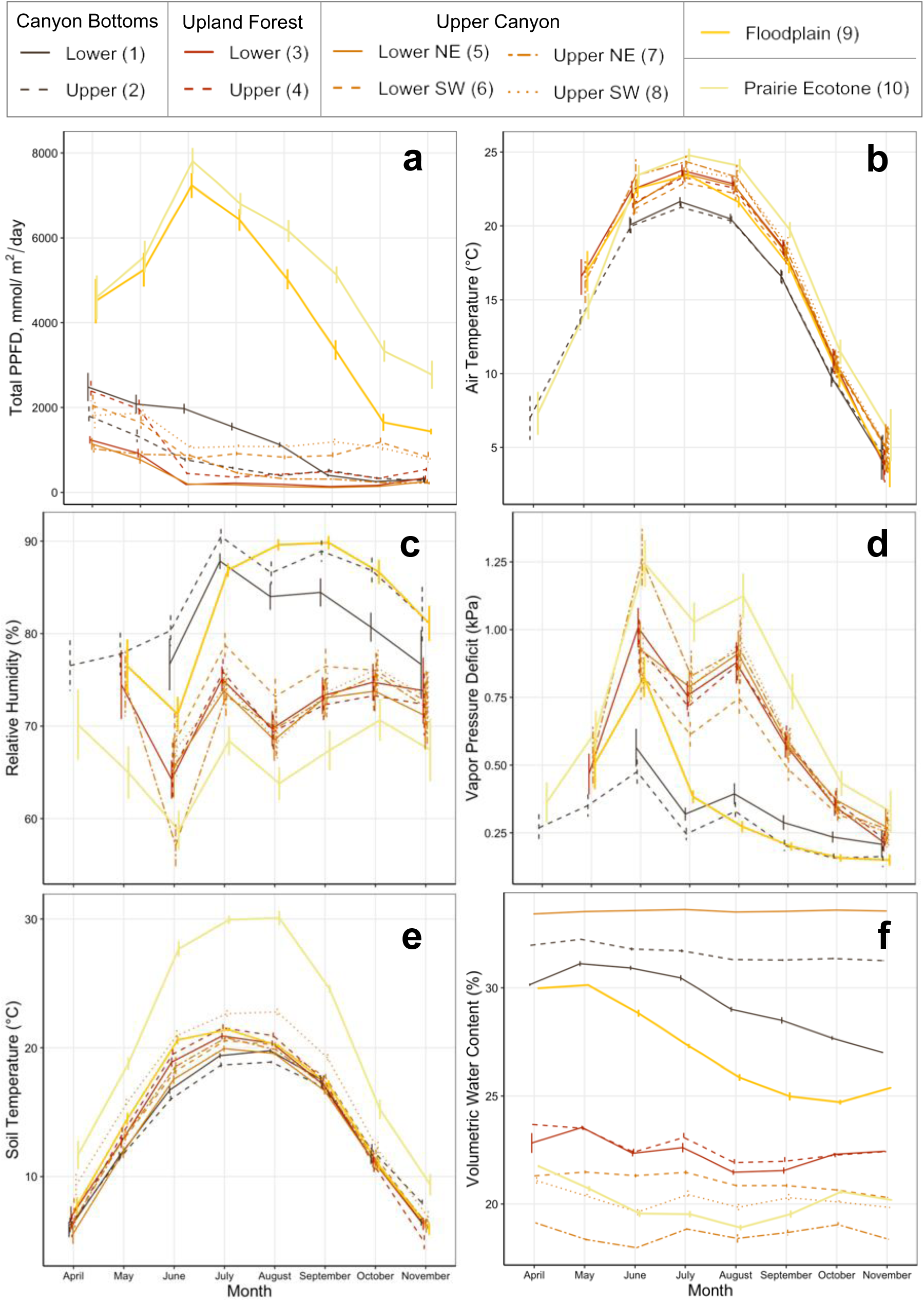
Habitat-related variation in microclimate in the Niobrara plot. Figures depict microclimate data collected at ten monitoring stations from April-November 2021. Monthly mean (± SE) values for (a) total daily photosynthetic photon flux density, (b) daily mean air temperature, (c) daily mean relative humidity, (d) daily mean vapor pressure deficit, (e) daily mean soil temperature, and (f) daily mean volumetric water content. Air temperature and relative humidity sensors were not deployed until 21 May 2021 for Stations 3, 7, and 9, and until 12 June 2021 for Stations 1, 4, 5, 6, and 8. Colors indicate habitats (Figure 1f), and different line types of the same color indicate data from stations within that habitat, with station numbers indicated in the legend and corresponding to Figure S1. In figure legend: NE = Northeast; SW = Southwest.

In multivariate analyses, habitats differed in their average microclimates (Figure S4a), with the first two principal components explaining 71.0% of the variation in microclimate. Overlap between the floodplain and canyon bottoms stations was driven by VWC and RH, and the upland forest and upper canyon had similar average microclimates. The prairie ecotone was distinguished from other habitats due to its high PPFD, air and soil temps, and VPD, and low VWC and RH (Figure S4a). The perMANOVA revealed that average microclimate varied significantly due to habitat type and the interaction between habitat and measurement month (habitat: *F*_4,40_ = 52.7, *R^2^* = 0.39, *p* = 0.001; habitat × month: *F*_35,40_ = 8.4, *R^2^* = 0.54, *p* = 0.001). Although habitats differed in microclimate variability (Figure S4b), the differences between them were not as pronounced as for the average microclimate (Figure S4a). Microclimate variability was more strongly driven by the month of measurement, with considerable overlap between habitats but clustering based on month (Figure S4b). While both habitat type (*F*_4,40_ = 11.2, *p* = 0.001) and the interaction between habitat type and measurement month (*F*_35,40_ = 9.4, *p* = 0.001) had significant effects on microclimate variability, the main effect of habitat type alone explained less variation (*R^2^* = 0.11) than the habitat-month interaction (*R^2^*= 0.80).

Variation in average microclimate was strongly influenced by variation in topography, with significant effects of topographic PC1 (*F*_1,69_ = 34.8, *R^2^* = 0.09, *p* = 0.001), PC2 (*F*_1,69_ = 16.8, *R^2^* = 0.04, *p* = 0.001), and PC3 (*F*_1,69_ = 55.0, *R^2^* = 0.14, *p* = 0.001), as well as month (*F*_7,69_ = 29.7, *R^2^* = 0.55, *p* = 0.001). While we also found microclimate variability to be significantly affected by topographically driven variation, less variance was explained by topographic PC1 (*F*_1,69_ = 3.8, *R^2^* = 0.01, *p* = 0.016), PC2 (*F*_1,69_ = 6.6, *R^2^* = 0.02, *p* = 0.002), and PC3 (*F*_1,69_ = 13.9, *R^2^* = 0.04, *p* = 0.001) compared to month (*F*_7,69_ = 34.6, *R^2^* = 0.72, *p* = 0.001).

Throughout the growing season, we found surface soil VWC (manually measured) to vary significantly among habitats (*F*_4,71_ = 2.9, *p* = 0.03) and sampling dates (*F*_5,326_ = 86.7, *p* < 0.001; Figure 3a), but there was no significant interaction between habitat and date (*F*_20,326_ = 1.3, *p* = 0.20), indicating differences in soil moisture between habitats remained consistent throughout the growing season. We found that the random effect of quadrat was significant (*p* < 0.001), indicating substantial spatial variation in soil moisture within habitats. The fixed effects of habitat and sampling date explained <25% of the variation in VWC (marginal *pR*^2^ = 0.24), whereas a high proportion of variation in VWC was cumulatively explained by habitat, date, and the random effect of quadrat (conditional *pR*^2^ = 0.87). The only pair of habitats that differed significantly in VWC in post-hoc comparisons was the floodplain and prairie ecotone (*p* = 0.01). Conversely, VWC differed significantly between most pairs of measurement time points (*p* ≤ 0.05), except for between early June (6/3/21) and mid-July (7/12/21; *p* = 0.24), early June and mid-October (10/10/21; *p* = 0.99), mid-July and mid-August (8/12/21; *p* = 0.18), and late July (7/23/21) and mid-October (*p* = 0.65). Although differences were not statistically significant, variation among habitats in the manually measured (greater spatial than temporal replication) VWC paralleled measurements from the microclimate stations (greater temporal than spatial replication) in that the floodplain and canyon bottoms had higher VWC than the upper canyon and prairie ecotone, creating a water availability gradient from the floodplain to prairie ecotone (Figure 3a).

**Figure 3.**
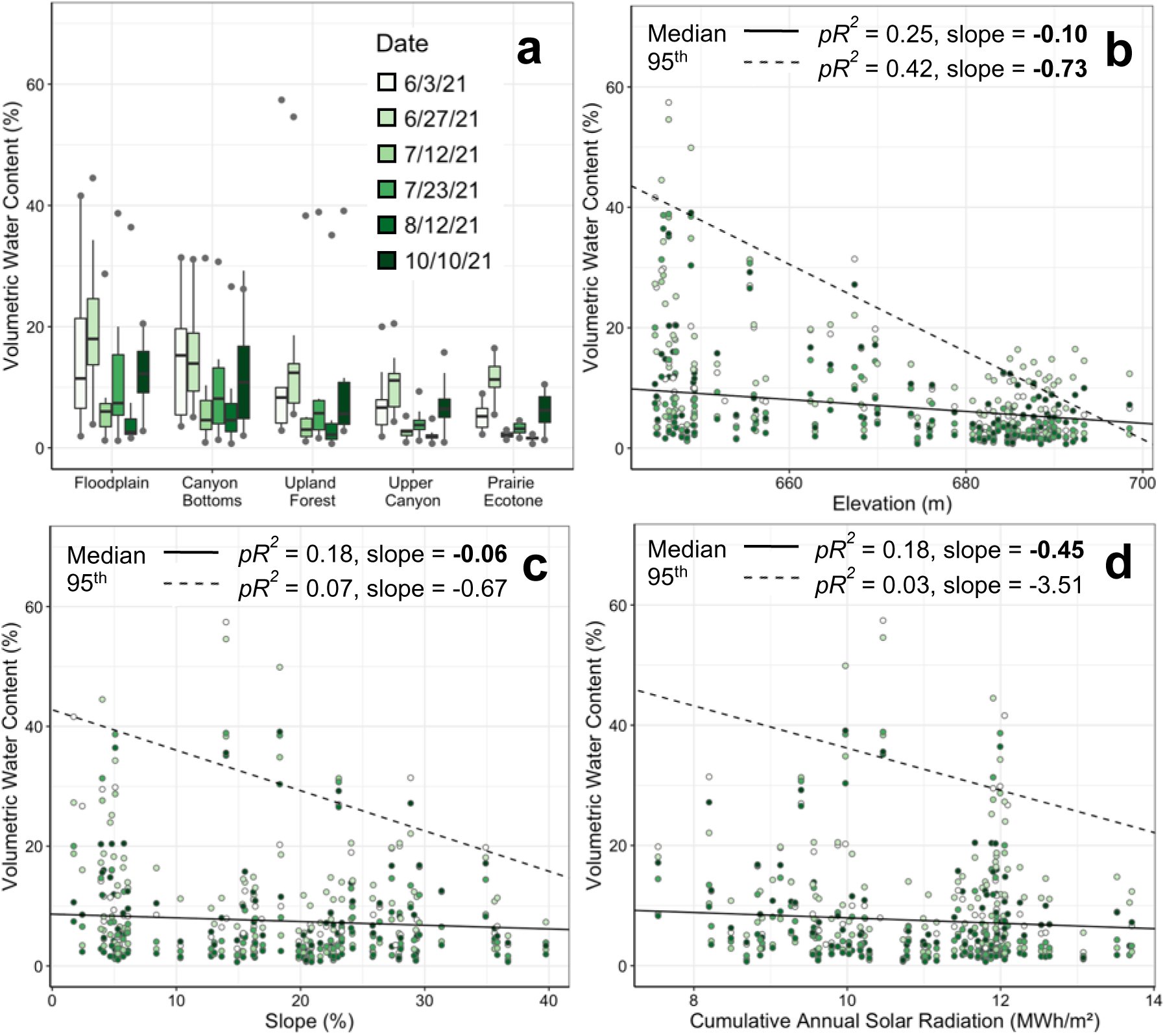
Variation in soil moisture across the growing season with respect to habitats and topographic variables showing. (a), variation in volumetric water content (VWC) in five habitats throughout the growing season. VWC was measured at six time points (white-dark green color ramp with earlier dates in lighter colors), and habitats are ordered by increasing elevation (Figure 1f). (b) – (d), scatterplots of VWC versus (b) elevation, (c) slope, and (d) cumulative annual solar radiation. In (a), boxes indicate the median (center line) and first and third quartiles; lower and upper whiskers indicate the first and third quartiles ±1.5 times the interquartile range; points indicate the minimum and maximum VWC. In (b) – (d), lines are predicted values for the median (solid line) and 95^th^ quantile (dashed line) relationships based on quantile regression models including an additive effect of date. *pR^2^* values in each panel are pseudo-*R^2^*, and bold font indicates quantile regression slopes that were significantly different from zero (*p* < 0.05).

Surface soil VWC (manually measured) also varied with topography throughout the growing season (Figure 3b-d). The relationship between VWC and elevation depended on VWC (Figure 3b); higher VWC values declined much more sharply with increasing elevation (95^th^ quantile *pR^2^* = 0.42, *F*_6,420_ = 60.7, *p* < 0.001) than lower values (median: *pR^2^* = 0.25, *F*_6,420_ = 54.7, *p* < 0.001), and although lower elevations displayed a range of VWC values, higher elevations displayed only low VWC. VWC declined with increasing slope (Figure 3c), more so for moderate VWC values (median *pR^2^* = 0.18, *F*_6,420_ = 58.2, *p* < 0.001) than for higher VWC values (95^th^ quantile *pR^2^* = 0.07, *F*_6,420_ = 1.1, *p* = 0.37). A similar pattern was observed for solar radiation (Figure 3d) in that VWC declined with increasing exposure for moderate VWC values (median *pR^2^* = 0.18, *F*_6,420_ = 50.0, *p* < 0.001), but not for higher VWC values (95^th^ quantile *pR^2^* = 0.03, *F*_6,420_ = 1.0, *p* = 0.42).

### 3.2 P2 Significant variation in forest structure, diversity, and composition among habitats and with respect to topography

Among trees, forest structure varied among habitats (Figure 4a-c; Table S5). Tree stem density differed significantly across habitats (Table S6), from a low of 25 (all habitats except canyon bottoms) to 4925 (floodplain) stems per ha, although this floodplain quadrat was an outlier with many small *Salix* spp. stems. Stem density was higher in the moister, lower-light habitats occurring at mid-elevations (canyon bottoms, upland forest, and upper canyon) compared to the more exposed, higher-light habitats (floodplain and prairie ecotone) (Figure 4a) and varied significantly between almost habitat pairs, except for the upland forest and upper canyon (*p* = 1.0). Total basal area also varied significantly across habitats (Table S6; Figure 4b), from 0.008 (prairie ecotone) to 68.70 (canyon bottoms) m^2^ per hectare. Basal area was higher in lower-exposure, mid-elevation habitats with moderate to high water availability (canyon bottoms and upland forest), with significant differences between many pairs of habitats [except canyon bottoms versus upland forest (*p* = 0.78) and upper canyon versus floodplain (*p* = 0.29)]. Total AGB ranged from 0.01 (prairie ecotone) to 482.26 (upland forest) Mg per ha, with significantly higher AGB in moister, lower-exposure habitats (upland forest and canyon bottoms) compared to the higher-exposure prairie ecotone (Table S6; Figure 4c). There were significant differences in AGB between most habitat pairs, except for the canyon bottoms and upland forest (*p* = 1.0) and the upper canyon and floodplain (*p* = 0.83).

**Figure 4.**
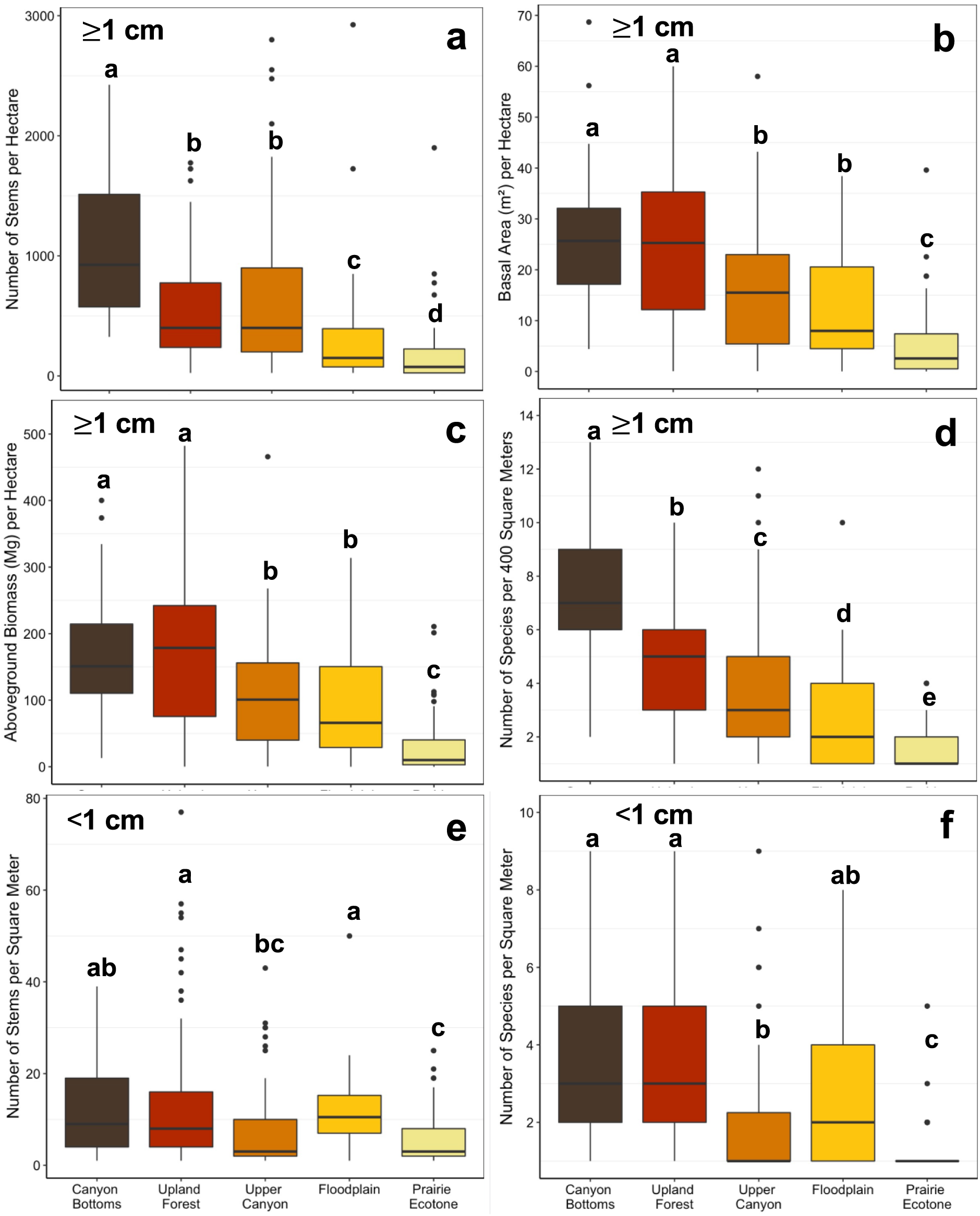
Variation in forest structure and diversity among five habitats in the Niobrara plot. Boxplots show habitat-related variation for tree (DBH ≥1 cm) (a) stem density per hectare, (b) basal area per hectare, (c) aboveground biomass per hectare, and (d) species richness per quadrat (400 m^2^), and for seedling (DBH <1 cm) (e) stem density per square meter and (f) species richness per square meter. Habitats are ordered by increasing light intensity and exposure from canyon bottoms to prairie ecotone (Figure 1f). Letters above the boxes indicate significant pairwise differences at the *α* = 0.05 level after correction for multiple comparisons using Tukey’s Honestly Significant Differences. Boxes indicate the median (center line) and first and third quartiles; lower and upper whiskers indicate the first and third quartiles ±1.5 times the interquartile range; points indicate outliers. One extreme outlier (a quadrat with 4925 stems/ha in the floodplain habitat) was excluded from (a) for better visualization of patterns; statistical analyses with and without this quadrat were consistent with each other. Means and standard errors are reported in Table S5.

Among trees, species richness varied significantly between habitats (Table S6), from 1 (all habitats except canyon bottoms) to 13 (canyon bottoms) species per quadrat (per 400 m^2^) (Figure 4d; Table S5). Species richness was higher in habitats with more complex forest structure that had cooler, moister microclimates (canyon bottoms) as compared to higher-light, hotter habitats (floodplain and prairie ecotone), and differed significantly between all pairs of habitats (all *p* < 0.05). Shannon’s diversity indices ranged from 0 (only one species present; all habitats except canyon bottoms) to 2.07 (canyon bottoms) per quadrat (per 400 m^2^), with significant differences occurring between habitats (Table S6). Diversity was higher in the moister, cooler canyon bottoms compared to the exposed prairie ecotone (Figure S5a), and diversity also differed significantly between all habitat pairs except upper canyon and floodplain (*p* = 0.25).

Seedling stem density also varied significantly between habitats (Table S6), from 1 (all habitats) to 77 (upland forest) stems per m^2^ (Figure 4e; Table S5). In contrast to the trees, variation in seedling stem density was less pronounced, and was higher in habitats with both higher light and higher water availability (Figure 4e). There were significant differences in seedling density between the canyon bottoms and prairie ecotone (*p* = 0.007), floodplain and prairie ecotone (*p* = 0.002), floodplain and upper canyon (*p* = 0.02), upland forest and prairie ecotone (*p* < 0.001), and upland forest and upper canyon (*p* = 0.006). For both seedling and tree size classes, the upland forest and canyon bottoms habitats had the greatest and second greatest number of species reaching their highest stem density in the Niobrara plot (Table S5, S7; number of species that achieve their greatest stem density by habitat: canyon bottoms = 10, upland forest = 11, upper canyon = 6, floodplain = 7, prairie ecotone = 3).

Seedling species richness also varied between habitats (Figure 4f; Table S5), ranging from 1 (all habitats) to 9 (canyon bottoms, upland forest, and upper canyon) species per m^2^. As with stem density, species richness of seedlings was higher in cooler, moister, and higher-light habitats (canyon bottoms, upland forest, and floodplain) than in higher elevation, exposed habitats. There were significant differences in seedling richness between the following habitats: canyon bottoms and prairie ecotone (*p* < 0.001), canyon bottoms and upper canyon (*p* = 0.001), floodplain and prairie ecotone (*p* < 0.001), upland forest and prairie ecotone (*p* < 0.001), and upland forest and upper canyon (*p* < 0.001). Shannon’s diversity indices for seedlings varied significantly among habitats (Table S6), ranging from 0 (all habitats) to 2.00 (canyon bottoms), and like richness, these values were higher in cooler, moister, and higher-light habitats (Figure S5b). Seedling diversity differed significantly between canyon bottoms and prairie ecotone (*p* < 0.001), canyon bottoms and upper canyon (*p* = 0.003), floodplain and prairie ecotone (*p* < 0.001), upland forest and prairie ecotone (*p* < 0.001), upper canyon and prairie ecotone (*p* = 0.04), and upland forest and upper canyon (*p* < 0.001).

Among trees, species composition varied significantly among habitats (*F*_4,334_ = 16.1, *R^2^* = 0.16, *p* = 0.001), and there was significantly greater within-habitat similarity in species composition than between-habitat similarity (*R*-value = 0.27, *p* = 0.001). Habitats differed in their variability in composition (*F*_4,334_ = 8.2, *p* < 0.001), with the least variable composition occurring in the canyon bottoms and the most variable in the floodplain (average distances of quadrats to the median, unitless; canyon bottoms: 0.997; upland forest: 1.184; upper canyon: 1.343; floodplain: 1.497; prairie ecotone: 1.263). There were significant pairwise differences in the variability in composition between the following habitats: floodplain and canyon bottoms (*p* = 0.001), prairie ecotone and canyon bottoms (*p* = 0.026), upper canyon and canyon bottoms (*p* < 0.001), and upland forest and floodplain (*p* = 0.003).

Based on analyses using continuous topographic variables, there were significant differences in forest structure, diversity, and composition, paralleling the findings based on categorical habitats. For tree stem density, among candidate models (Table S4), the most-supported model (75.6% of Akaike weight) included the three two-way interactions between elevation, eastness, and slope, and explained 69% of variation in stem density (Table S8). There was a significant main effect of elevation (*p* < 0.001), as well as significant interactions between elevation and slope (*p* < 0.001) and eastness and slope (*p* = 0.003) (Table S9). The positive effects of steeper slope on stem density were greater at higher elevations for both east and west aspects and, on western aspects, this was also the case for lower elevations (Figure 5a). While low-density quadrats never occurred on steeper slopes, quadrats with low to moderately steep slopes had a wide range of stem densities depending on elevation and aspect (Figure 5a). For total basal area, the most-supported model (48.3% of weight) included all two-way interactions between elevation, northness, and slope and explained 56% of variation in basal area (Table S8). There was a significant main effect of elevation (*p* = 0.010), and significant interactions between elevation and slope (*p* = 0.018) and northness and slope (*p* = 0.007; Table S9). Like stem density, we found a positive effect of steeper slope on basal area, though the overall relationship was more variable, and the effect of slope was generally stronger at higher elevations for south-facing aspects (Figure 5b). For total AGB, the most-supported model (22.7% of weight) included two-way interactions between elevation, northness, and slope, as with basal area, explaining 52% of variation in AGB (Table S8). However, the weight for the second-best model including only the interaction between slope and northness was almost equivalent (Akaike weight = 22.1%, adjusted *R*^2^ = 0.52; Table S8). The importance of the slope-northness interaction in the best model (*p* = 0.027) is underscored by the marginally significant interaction between elevation and slope (*p* = 0.051) and non-significant interaction between elevation and northness (*p* = 0.82), although the main effect of elevation was also significant (*p* = 0.024; Table S9). The slope-northness interaction was strongest in quadrats with northern aspects (gray points) and higher elevations, and more variable in quadrats with southern aspects (Figure 5c).

**Figure 5.**
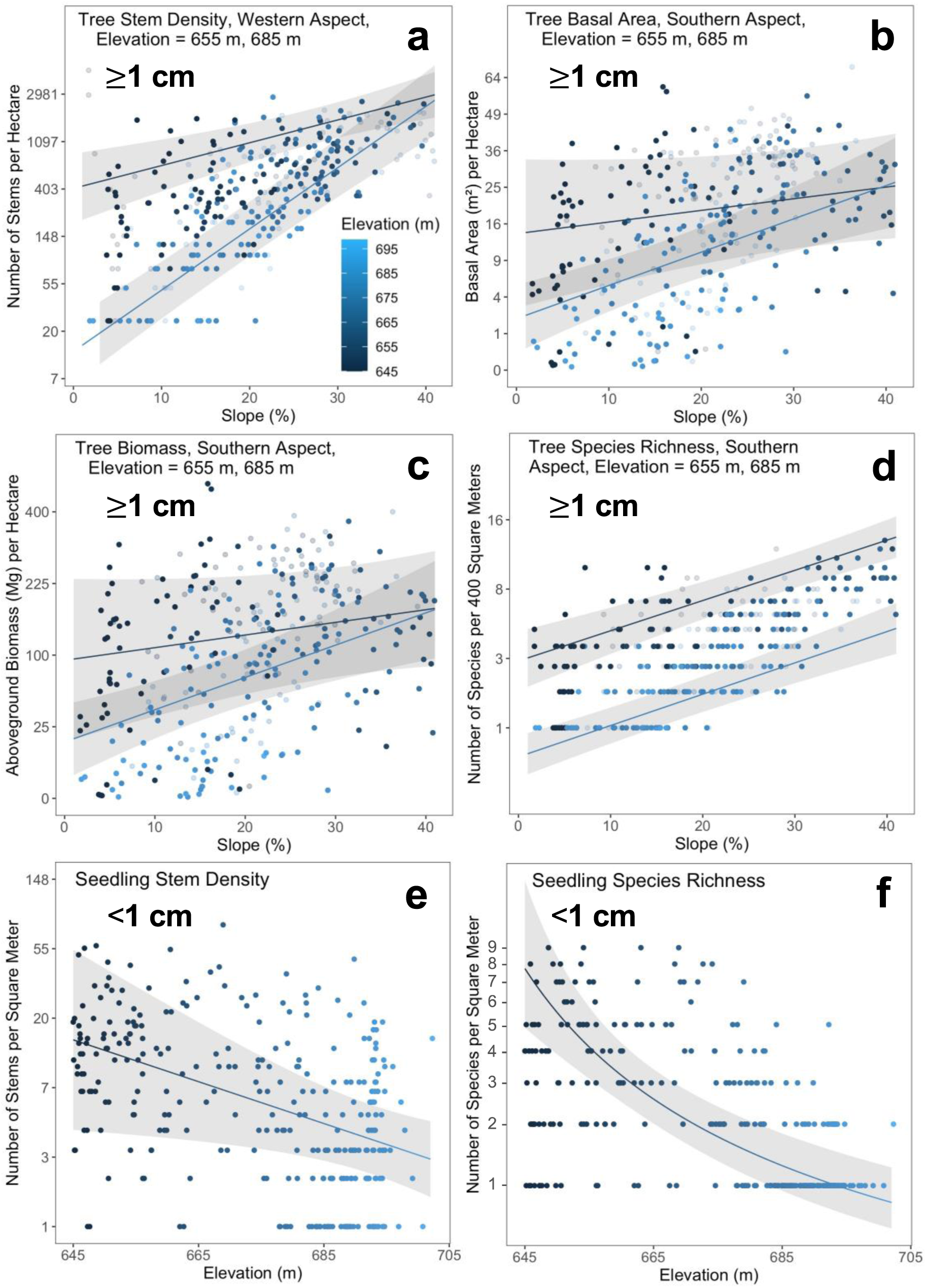
Continuous variation in forest structure and diversity with respect to topographic gradients in the Niobrara plot. Scatterplots (points) with predictions (lines and 95% confidence ribbons) based on the best candidate general additive models showing variation in tree (DBH ≥1 cm) (a) stem density per hectare, (b) basal area per hectare, (c) aboveground biomass per hectare, (d) species richness per quadrat, and seedling (ground diameter <1 cm) (e) stem density per square meter and (f) species richness per square meter versus slope (a-d) and elevation (e-f). All response variables and elevation have been back transformed, and the best candidate models for each variable are listed in Table S8. For (a)-(d), predictions for low and high elevations were generated by setting elevation equal to the first and third quartiles (655 and 685 m), indicated by the dark and light blue fitted lines, respectively, and specifying a (a) western (eastness = −1) or (b)-(d) southern (northness = −1) aspect, respectively. For (e) and (f), predictions were generated across all elevations. Each point represents data from a quadrat (a-d) or seedling subplot within a quadrat (e,f), color-coded based on its elevation as indicated by the color bar in the in-figure legend. Points in the blue gradient (a)-(d) correspond to quadrats with (a) western-facing (eastness = −1-0) or (b)-(d) southern-facing (northness = −1-0) aspects, whereas grayed out points have (a) eastern-facing (eastness = 0-1) or (b)-(d) northern-facing (northness = 0-1) aspects.

For tree species richness, the most-supported model (44.2% of weight) included all two-way interactions between elevation, northness, and slope, capturing 76% of the variation in richness (Table S8). The main effects of elevation (*p* < 0.001), northness (*p* = 0.045), and slope (*p* < 0.001), and the interaction between elevation and northness (*p* = 0.003), were significant (Table S9). Species richness increased considerably on steeper slopes across both north and south aspects, and the intercept for species richness was higher for lower elevations on south-facing aspects, indicating higher richness occurred at lower elevations (Figure 5d).

Among trees, there was strong variation in species composition between quadrats along the topographic gradient (Figure 6f; lowest stress value: 0.143; non-metric *R*^2^ = 0.98; linear *R*^2^ = 0.87). While continuous topographic variation significantly influenced tree species composition (full model: *R*^2^ = 0.16, *p* = 0.001), there were strong differences in the variance in composition explained by each topographic variable, and only the effects of elevation, solar radiation, and slope were significant. Elevation explained the highest amount of variation (*R*^2^ = 0.08, *p* = 0.001), followed by slope (*R*^2^ = 0.03, *p* = 0.001), solar radiation (*R*^2^ = 0.002, *p* = 0.03), eastness (*R*^2^ = 0.001, *p* = 0.09), and northness (*R*^2^ < 0.001, *p* = 0.56).

**Figure 6.**
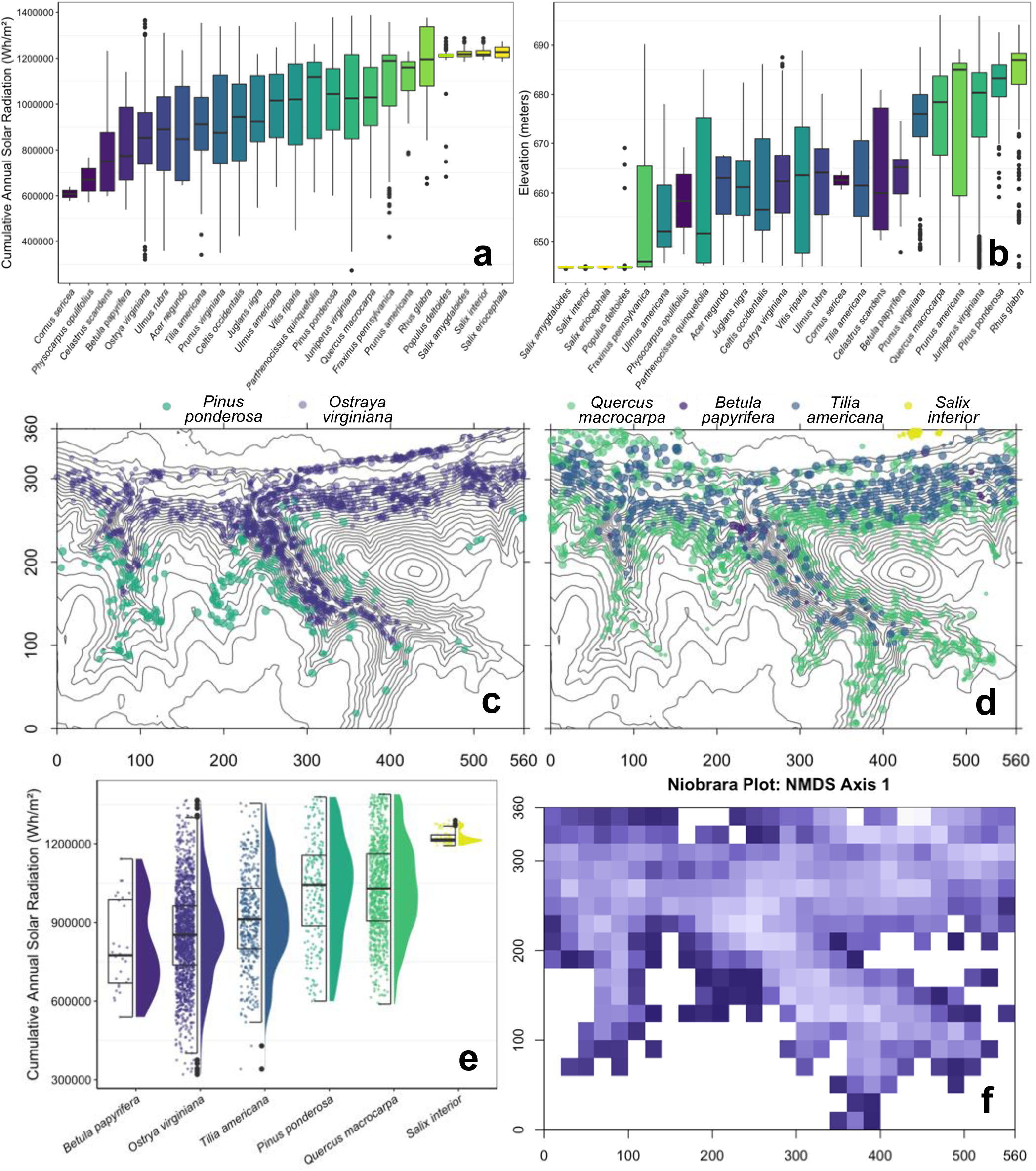
Topographic niches, species distributions, and variation in community composition in the Niobrara plot. (a) – (b) Boxplots showing the a) solar radiation niches and b) elevation niches for woody species with >1 individuals in the plot, ordered by median value for each environmental variable, and color-coded by increasing median cumulative annual solar radiation. Boxes indicate the median (center line) and first and third quartiles; lower and upper whiskers indicate the first and third quartiles ± 1.5 times the interquartile range; points indicate outliers. (c) – (d) Topographic maps of the Niobrara plot with 5-m elevation contour lines, showing species distributions (color-coded by median solar radiation) for c) *Pinus ponderosa* and *Ostrya virginiana*, and d) *Quercus macrocarpa*, *Betula papyrifera*, *Tilia americana*, and *Salix interior*. (e) Combined boxplot and violin plot showing the solar radiation niches and density of individuals within the niche for the six species whose distributions are mapped in (c) and (d) (color-coded by median solar radiation), with boxes indicating the same values described in (b). (f) Map of the scores from the first axis of a non-metric multidimensional scaling of the Niobrara plot community composition. A Cao dissimilarity matrix of between-quadrat differences in community composition was used, and each colored square represents a 20×20 m quadrat, with similarly shaded squares having more similar species composition, and white squares representing quadrats with no woody stems ≥ 1 cm diameter at breast height.

Among seedlings, continuous topographic variation also explained considerable variation in structure and diversity. The most-supported model for seedling stem density (8.2% of weight) only included the main effect of elevation (*p* < 0.001; Table S9), explaining 30% of variation in seedling stem density. In contrast to results for tree stem density, seedling stem density generally decreased with increasing elevation, although this effect was variable (Figure 5e). For species richness of seedlings, the top model (18% of weight) also only included the main effect of elevation (*p* < 0.001; Table S9) and captured 49% of the variation in seedling species richness. Seedling species richness decreased with increasing elevation (Figure 5f), although subplots at lower elevations showed greater variability in richness compared to the consistently low richness at higher elevations.

### 3.3 P3 Topographic niches of species structure forest diversity and composition

Topography strongly shaped species’ distributions across the plot (Figure 6a-e). Species occupied well defined solar radiation and elevation niches (Figure 6a, b). Some species exhibited very narrow topographic niches (*e.g.*, *Salix interior*, *B. papyrifera*), indicating strong habitat affinities (Figure 6a, b; clustered distribution of *S. interior* along the floodplain and *B. papyrifera* near groundwater springs in the canyon bottoms in Figure 6d). Conversely, other species exhibited wider topographic niches. For example, *Ostrya virginiana* was more concentrated in lower-light environments but is widespread throughout the canyon bottoms and midstory of upland forest (Figure 6a, c, e), and *Q. macrocarpa* is distributed widely in higher-light environments from the floodplain to prairie ecotone (Figure 6a, d, e).

### 3.4 P4 Topographically driven microclimatic variation influences forest composition

Species composition of trees varied significantly with microclimate and its variability. Based on perMANOVA, VPD had a significant effect on species composition of trees and uniquely explained a substantial amount of its variation (VPD: *F*_1,5_ = 2.4, *R^2^* = 0.25, *p* = 0.03). Although the effects of soil temperature (*F*_1,5_ = 1.7, *R^2^* = 0.18, *p* = 0.14) and moisture (*F*_1,5_ = 0.9, *R^2^* = 0.10, *p* = 0.51) were not statistically significant, they each had considerable, unique explanatory power, and including the co-explained variance, together they explained 52.2% of the variation in composition. The results from the MRM were similar, although the effect of mean VPD was only marginally significant (*p* = 0.058), and the effects of soil temperature (*p* = 0.28) and VWC (*p* = 0.13) were not statistically significant. Based on the perMANOVA, the effects of variability in VPD and soil temperature were statistically significant, and both explained considerable variation in species composition (VPD: *F*_1,5_ = 2.4, *R^2^* = 0.25, *p* = 0.05; soil temperature: *F*_1,5_ = 2.9, *R^2^* = 0.30, *p* = 0.006) compared to VWC (*F*_1,5_ = 0.3, *R^2^* = 0.04, *p* = 0.92). Together, variability in VPD and soil temperature and moisture explained a total of 59.0% of the variation in composition. However, based on the MRM, none of these effects was statistically significant (all *p* > 0.40).

## 4. Discussion

Although biomes are defined at large spatial scales, shifts in their boundaries arise through the aggregated responses of individuals to their local conditions, including microclimate, along biome transition zones (Oliveras & Malhi, 2016). To predict the responses of vegetation to climate change and potential for biome shifts, it is essential to understand microclimatic variation, the factors influencing it, and how vegetation responds to it, as we have done here for this semi-arid forest at the forest-grassland transition zone in North America. To our knowledge, our study is the first to have done so by integrating forest inventory data on size classes from seedlings to adults with detailed topographic and locally measured microclimatic data along the environmental gradients across this transition zone.

These gradients were characterized by sharp variation in conditions affecting desiccation, driven by differences in exposure and water availability. The magnitude of the topographically driven microclimate variation in the Niobrara plot across the forest-grassland transition zone is on the order of macroclimate variation at larger spatial and temporal scales. The 2.2 °C mean air temperature difference along the 59-m gradient from the floodplain to the prairie ecotone in the Niobrara plot is equivalent to the mean annual temperature change across approximately three degrees of latitude in North America (La Sorte et al., 2014). Based on remotely estimated soil moisture averaged across the 2021 growing season from the NASA Soil Moisture Active Passive product (NASA GSFC, 2020), the 10.2% mean difference in soil moisture across the elevational gradient in the Niobrara plot approximately corresponds to the average difference in soil moisture across approximately 2.5 degrees of longitude to the east (wetter) along the east-west aridity gradient in the North American Great Plains. In a seasonal context, the magnitudes of these habitat-related microclimatic differences are equivalent to the change in daily regional air temperature between late June and the hottest day of the growing season in late July (based on daily averages from 1981-2010; U.S. Climate Data, 2022), and in soil moisture from early May to mid-June (NASA GSFC, 2020).

Topographically driven microclimatic variation strongly affected forest structure, diversity, and composition in the Niobrara plot across the forest-grassland transition zone. While heterogeneity in microclimatic conditions driven by topography is critical in structuring plant communities along more dramatic alpine elevational gradients (Opedal et al., 2015; Rae et al., 2006), we found it was also a key driver of composition along the much shorter elevational gradient in the semi-arid Niobrara forests. Woody species exhibited well-defined solar radiation and elevational niches, causing high species turnover and differences in forest composition between habitats with distinct topography and microclimate. More structurally complex and diverse stands with predominantly broad-leaved, mesic forest species (*e.g.*, *T. americana*, *Ulmus rubra*) were found on steeper slopes and in less exposed, cooler, moister habitats. Conversely, less structurally complex, and less diverse stands with more widespread drought-tolerant species (*e.g*., *Q. macrocarpa*, *P. ponderosa*) were found in more exposed, hotter habitats at higher elevations along the transition zone. This zone abutted the hottest, driest conditions of the regionally dominant grassland biome, in which only a few woody shrub species occurred (*e.g.*, *Rhus glabra*, *Rosa arkansana*). The shelter from the regional macroclimate that is provided by the Niobrara River valley and its associated canyons, with their consistently lower VPD, forms microclimate refugia that allow tree species from diverse provenances and with varied physiological tolerances to persist, many at their geographic range edges. Our study suggests that the semi-arid Niobrara forests both depend on and moderate these microclimate refugia and fills a critical gap by providing new insight on the relative importance of key topographic and microclimatic variables driving the biome transition from forest to grassland and structuring this forest, knowledge that is crucial in quantifying the effects of climate change on these zones.

### 4.1 Microclimate effects on boundary forest communities

Local microclimate conditions can differ considerably from the macroclimate, and this spatial heterogeneity in environmental conditions will differentially affect species’ abilities to recruit and survive (Blonder et al., 2018; Potter et al., 2013). For tree seedlings, the effects of climate warming can be buffered by the overstory reducing temperatures and exposure in the understory and creating more favorable conditions for establishment (De Frenne et al., 2013; Scherrer & Körner, 2010), particularly in semi-arid systems. The lower understory light availability, cooler temperatures, and higher soil moisture we documented in the canyon bottoms and upland forest are evidence of such buffering. In the Niobrara plot, this buffering arises from both the microclimate refugia created by the physical environment, as well as the biotic effects of the forest overstory, the presence of which in turn depends on these refugia. Such microclimate refugia may reduce the effects of macroclimatic changes (Dobrowski, 2011; Hampe & Jump, 2011), but this idea is not well tested across biomes (Carnicer et al., 2021; Hacket-Pain & Friend, 2017) and warrants investigation.

Soil moisture gradients are known to structure forest-grassland transition zones (Pool et al., 1918; Terra et al., 2018) and have been estimated using soil and topographic variables in hydrological models (Neupane & Kumar, 2015; Sloan & Moore, 1984). Yet, difficulty in monitoring water availability gradients at fine spatiotemporal scales has limited their empirical quantification along vegetation transitions (Yeakley et al., 1998). Previous studies have documented how forest structure and diversity vary with topography, assuming that topography is a direct proxy for water availability as the principle driver (Jucker et al., 2018; Méndez-Toribio et al., 2016; Muscarella et al., 2020; Valencia et al., 2004; Vieira et al., 2004), but few studies (Kupers et al., 2019; Ma et al., 2010; Russo et al., 2010) have validated these assumptions with detailed soil moisture data, as we did in this study. Our monitoring revealed that temporal variation in soil moisture within habitats can be as high as variation between habitats, possibly owing to the well-drained soils of our study site. Soil moisture exhibited great variability at lower elevations but was consistently lower at higher elevations along the forest-grassland transition zone, where the well-drained soils favor grasses, forbs, and drought-tolerant woody species over mesic forest components (Kantak, 1995). The narrow canyon bottoms were more buffered from this temporal variation due to their lower elevation and the influence of groundwater seeps, which moderated soil moisture and temperature, allowing for local persistence of species characteristic of more mesic forests (Kaul et al., 1988; Tolstead, 1942a).

Semi-arid ecosystems, such as those in Great Plains of North America are challenging for tree growth owing not only to insufficient rainfall during the growing season, but also to its highly pulsed nature (Woods et al., 2014; Muñoz-Rojas et al., 2016; Kaul et al., 1988). Our rainfall data is consistent with these observations in that most growing season rain fell during a few, high-precipitation storms. Due to climate change, such extreme precipitation events may be increasing in frequency and intensity (Bishop et al., 2021; Singh et al., 2013). In the well-drained, sandy soils characteristic of the Niobrara plot and surrounding region, extreme precipitation events may render soil water less available to plants (Muñoz-Rojas et al. 2016), as less infiltration occurs, resulting in more rainfall being lost to runoff and increasing soil erosion (Shrestha et al., 2021; Wu et al., 2018). One exceptionally large rainstorm in mid-July (Figure S3a) caused significant erosion and sedimentation in the main canyon (B. McNichol, personal observation). However, soil moisture only briefly increased before returning to drier conditions in all habitats except for the canyon bottoms (Figure 2f), suggesting benefits of this rainfall pulse for trees may have been short-lived.

Tree species differ in sensitivity to water limitation and the degree to which their growth is buffered by access to groundwater (Chitra-Tarak et al., 2018, 2021; Costa et al., 2022; Dawson & Ehleringer, 1991; Pettit & Froend, 2018). Such differences are seen even between drought-tolerant species, affecting their long-term growth responses to aridity (Aus Der Au et al., 2018). In the Niobrara forests, rainfall has been hypothesized to be less critical for growth and survival of woody species compared to groundwater access (Kaul et al., 1988; Szilagyi et al., 2003). In a nearby plantation in the Nebraska Sandhills, seasonal differences were observed in soil water uptake depth by *J. virginiana* and *P. ponderosa*, with both species using considerably deeper water (> 0.9 m depth) during the driest part of the growing season (Eggemeyer et al., 2008). However, many woody species in the Niobrara forests are likely to be considerably less tolerant of water limitation than either of these conifers (*e.g., T. americana*; Abrams et al., 1998), which raises questions about future changes in forest composition driven by shifts in habitat and geographic distributions of the many species at the edges of their ranges in the Niobrara region. This is particularly true considering the increasing macroclimate aridity, concentration of rainfall in large storm events, and potential reductions in tree access to groundwater caused by increasing regional demand for agricultural irrigation, which is largely supplied by groundwater (McGuire, 2011).

### 4.2 Role of topographic gradients in shaping boundary forest structure, diversity, and composition

Forest structure, diversity, and composition corresponded strongly to topographic and microclimatic gradients in the Niobrara forests. We observed more structurally complex stands on lower-elevation, steep slopes in moister, cooler habitats, with higher tree stem density occurring on west-facing slopes, and higher tree basal area and AGB occurring on south-facing slopes. Because they are often positively associated (Slik et al., 2010), we expected similar patterns of stem density and basal area with respect to topographic gradients in the Niobrara plot, and we found both to be higher in mesic, lower-exposure areas. However, west-facing aspects promoted higher stem density, paralleling findings in North American deciduous forest (Fekedulegn et al., 2003), whereas south-facing aspects facilitated higher basal area, which was observed in semi-arid Himalayan forest (Måren et al., 2015). Higher basal area has also been associated with mesic lower-elevation swales, in North American deciduous forest (Smith et al., 2017) and in several forest types in northwestern South America (Álvarez-Dávila et al., 2017).

Spatial variation in basal area and AGB are also often positively correlated (Baraloto et al., 2011; Malhi et al., 2006). Although we found higher basal area on steeper slopes in the sheltered canyons (Figure 5b), the highest AGB occurred on more moderate upland forest slopes, likely driven by the high wood density of *Q. macrocarpa* (Miles & Smith, 2009), which is much more abundant there (Table S7). Woody primary productivity may be more strongly limited by soil moisture compared to air temperature (Helcoski et al., 2019), especially in water-limited, semi-arid forest ecosystems (Adams et al., 2014; Andrews et al., 2020). Our results underscore the roles of elevation and aspect in creating desiccation gradients that limit woody productivity (Adams et al., 2014; Helcoski et al., 2019) and drive the transition from forest to grassland biome. Thus, our findings are consistent with those from other forest types, emphasizing the general importance of water availability in increasing the structural complexity of forests.

Similar to patterns of basal area and AGB, we documented more diverse stands on lower-elevation, steeper, south-facing slopes in the moister canyon bottoms and upland forest. Under these topographic and microclimatic conditions, increased water availability and heterogenous light conditions likely increased richness by facilitating higher co-occurrence of species with differing requirements (Álvarez-Dávila et al., 2017; Lusk, 2019; Peterson & Reich, 2008). Like trees, seedling stem density and richness were higher in moister, lower-elevations habitats, but specifically those that also had higher understory light, which is critical for regeneration (Jin et al., 2017). Our evidence of topographic niche partitioning parallels the findings of topographic niches of tree species in other forest types (Pulla et al., 2017; Valencia et al., 2004; Harms et al. 2001; Duque et al., 2002). Trees of woody species in the Niobrara plot showed preferences for relatively more or less exposed habitats, reflected in their distributions along the elevation and total annual solar radiation gradients. The willow species (*Salix amygdaloides*, *S. eriocephala*, *S. interior*), small-stature trees occurring at high stem densities in the floodplain, were the only species found exclusively on a single habitat. While differences between topographic distributions of trees and seedlings may signal future changes in the structure and diversity of this forest and the forest-grassland transition zone, data on growth, survival, and recruitment is required to evaluate this hypothesis.

## 5. Conclusions

Our study provides novel insights on drivers of variation in forest structure, diversity, and composition in an ecologically important boundary forest at the forest-grassland transition zone in the North American Great Plains. Sharp topographic gradients and groundwater seeps strongly influenced microclimatic variation across small spatial scales, creating microclimate refugia supporting forests within the regional semi-arid macroclimate in which the grassland biome predominates. The topographically driven variation in microclimate corresponded with changes in forest composition, promoting co-existence of species with distinct topographic niches linked to their diverse geographic provenances and physiological tolerances, and driving the biome transition to grassland at higher elevations with increased exposure. As in past eras of dramatic climate change, we speculate that microclimate refugia, such as those in the Niobrara, will be critical in maintaining diversity by allowing forests to persist (*e.g.*, Bush, 2017; da Rocha, 2019; Haffer 1969). The refugia may ameliorate tree stress and mortality associated with larger-scale changes in macroclimate, increasing resilience by diverting these systems from climate change-driven tipping points that precipitate complete forest loss. Because microclimate variation driving the forest-grassland transition zone recapitulates the trajectory of climate change, investigations of the dynamics in these boundary forests may provide early indicators for how species in the core parts of forested biomes will respond to direct and indirect effects of climate change. This is particularly true in the Niobrara region, where most woody species and forest-dependent species are at the distributional edges of their habitat, geographic, and biome ranges.

## Supporting information

Electronic Supplementary Materials

## Acknowledgements

This study is dedicated to the memory of Robert Bruce Kaul (1935-2019), lead author of the *Flora of Nebraska* (Kaul et al. 2011), Curator of Botany at the University of Nebraska State Museum, colleague, and friend. His exceptional insights and botanical studies on the Niobrara region inspired the Niobrara plot and this research. We thank Sakia Fields, Jennifer Foster, Joseph Eastman, Townson Lemansky, Jack Luth, Brittni McGuire, Susana Moyer, Ashley Newman, Cooper Steffen, Patrick Wilson, and Jessica Shue for assistance with the first census of the Niobrara plot. We thank Drew Tyre (University of Nebraska–Lincoln, UNL) for advice on statistical analyses. We thank Kristina Anderson-Teixeira (Smithsonian Conservation Biology Institute) for providing PRISM monthly climate data (NSF #DEB-1353301) for the Niobrara plot. We acknowledge The Nature Conservancy for permission to install the Niobrara plot on the Niobrara Valley Preserve and for providing logistical support of this research. We thank Chad Bladow (The Nature Conservancy) for his assistance and support. We thank the UNL Department of Civil Engineering and School of Natural Resources for use of surveying equipment. This research was supported by funding from Jacqueline B. Mars, the Smithsonian ForestGEO program, the National Science Foundation AccelNet Program (OISE 2020424), the American Philosophical Society, the Botanical Society of America, the Nebraska Native Plant Society, the UNL Center for Plant Science Innovation, the UNL School of Biological Sciences Dr. John F. Davidson Memorial Fund, a UNL Faculty Seed Grant, the UNL Undergraduate Creative Activities and Research Experience program, and the Cabela’s Apprenticeship Program.

## Author Contributions

BHM and SER conceived of and designed the study, collected the data, conducted statistical analyses, and wrote the manuscript. RW derived the topographic variables from the DEM, assisted with making figures, and helped write the manuscript. AH and CH secured permission from The Nature Conservancy to establish the Niobrara plot on the Niobrara Valley Preserve and helped write the manuscript. SMM was involved in the establishment of the Niobrara plot and helped write the manuscript.

## Data Availability Statement

The Niobrara forest inventory plot data and microclimate data are available at: https://forestgeo.si.edu/explore-data/niobrara-termsconditionsrequest-forms. The digital elevation model for Brown County, Nebraska, is available at: https://www.sciencebase.gov/catalog/item/543e6b86e4b0fd76af69cf4c.

