## Supplementary material for "Topographically driven microclimatic gradients shape patterns of forest structure, diversity, and composition at a forest-grassland transition zone": Electronic Supplementary Materials

**Appendix S1: Supplementary Methods**

*Derivation of habitat types*

Following previous studies (*e.g.*, Kenfack et al., 2014; Valencia et al., 2004), we defined five categorical topographic habitats using the five topographic variables. To capture the covariation among these variables, we conducted principal components analysis (PCA) using scaled data as implemented in the *prcomp* function in the 'stats' package (R Core Team 2020). The first principal component (PC1; 42.7% of variation explained) was correlated with variation in slope, solar radiation, and elevation, and PC2 (22.6% of variation explained) was correlated with aspect (Figure S2a; Table S2). Habitats were defined based on the quadrats' scores for PC1 and PC2 using the following cutoffs (with habitats listed in order of increasing light intensity and exposure): 1) canyon bottoms: PC1 > 1.00, PC2 < 0.20; 2) upland forest: PC1 ≥ -0.20, PC2 ≥ 0.20; 3) upper canyon: -0.98 ≤ PC1 ≤ 1.00, PC2 < 0.20; 4) floodplain: -0.98 ≤ PC1 < -0.20, -0.72 ≤ PC2 ≤ 1.81; 5) prairie ecotone: PC1 < -0.98, -1.40 ≤ PC2 ≤ 1.62. Due to some areas of the plot having considerable overlap in PCs driven by different variables (*e.g.*, floodplain and prairie ecotone quadrats are high-exposure environments, but occur at the lowest and highest elevations, respectively), we manually reassigned 74 quadrats (14.6%) that had been assigned to illogical habitats based on their positions along the topographic gradients. The distribution of quadrats assigned to each habitat across the Niobrara plot is shown in Figure 1f (canyon bottoms: 2.04 ha; upland forest: 5.56 ha; upper canyon: 3.56 ha; floodplain: 1.48 ha; prairie ecotone: 7.52 ha). Habitats varied significantly in topographic variables (Figure S2a), indicating that our habitat definitions were reasonable, as supported by a permutational multivariate analysis of variance (perMANOVA) of the Bray-Curtis distances between quadrats in the scaled topographic variables as a function of habitat ($F_{4,334} = 117.6$, $R^2 = 0.58$, $p = 0.001$) implemented with the *adonis2* function in the 'vegan' package (Oksanen et al., 2020).

**Table S1. Taxonomy, growth form, size class, and habitat associations for the 37 woody species (27 reaching diameter at breast height (DBH) ≥1 cm) in the 20.2 ha Niobrara plot in semi-arid forest on the south side of the Niobrara River, Nebraska, USA.** Geographic range descriptions are qualitative, based on visual inspections of the distributions documented from aggregated journals and periodicals, monographs, and herbarium records by the Biota of North America Program. Size classes for species were defined as follows: Adult (A) = >5 cm DBH; Sapling (Sap) = ≥1-5 cm DBH; Seedling (Sdlg) = <1 cm diameter at ground level. Determinations of stem density (number of stems in the Niobrara plot/ha) and the habitat of greatest stem density only includes the individuals included in the full plot census (adults and saplings), and densities are standardized based on habitat area. A "—" for stem density indicates species that only had seedlings in the Niobrara plot.

| **Species (Family)** | **Growth form** | **Geographic range** | **Size class** | **Stem density** | **Habitat of greatest stem density** |
|---|---|---|---|---:|---|
| *Acer negundo* (Sapindaceae) | Canopy tree | Widespread | A Sap | 0.29 | Canyon bottoms |
| *Amorpha canescens* (Fabaceae) | Shrub | Central | Sdlg | — | Upland forest |
| *Betula papyrifera* (Betulaceae) | Canopy tree | Boreal | A Sap Sdlg | 2.06 | Canyon bottoms |
| *Celastrus scandens* (Celastraceae) | Liana | Eastern | A Sap Sdlg | 0.59 | Canyon bottoms |
| *Celtis occidentalis* (Cannabaceae) | Canopy tree | Eastern | A Sap Sdlg | 20.43 | Upland forest |
| *Cornus sericea* (Cornaceae) | Shrub | Widespread | Sap | 0.15 | Canyon bottoms |
| *Dalea villosa* (Fabaceae) | Shrub | Central | Sdlg | — | Prairie ecotone |
| *Fraxinus pennsylvanica* (Oleaceae) | Canopy tree | Widespread | A Sap Sdlg | 30.24 | Floodplain |
| *Juglans nigra* (Juglandaceae) | Canopy tree | Eastern | A Sap Sdlg | 1.99 | Upland forest |
| *Juniperus virginiana* (Cupressaceae) | Canopy tree | Eastern | A Sap Sdlg | 208.70 | Upper canyon |

| | | | | | |
|---|---|---|---|---|---|
| *Lonicera dioica* (Caprifoliaceae) | Liana | Eastern | Sdlg | — | Canyon bottoms |
| *Morus alba* (Moraceae) | Canopy tree | Non-native | A | 0.07 | Canyon bottoms |
| *Ostrya virginiana* (Betulaceae) | Midstory tree | Eastern | A Sap Sdlg | 115.71 | Upland forest |
| *Parthenocissus quinquefolia* (Vitaceae) | Liana | Eastern | A Sap | 1.77 | Floodplain |
| *Physocarpus opulifolius* (Rosaceae) | Shrub | Eastern | Sap Sdlg | 0.15 | Canyon bottoms & Upland forest |
| *Pinus ponderosa* (Pinaceae) | Canopy tree | Western | A Sap Sdlg | 16.15 | Upper canyon |
| *Populus deltoides* (Salicaceae) | Canopy tree | Widespread | A Sap Sdlg | 13.86 | Floodplain |
| *Prunus americana* (Rosaceae) | Understory tree | Widespread | Sap Sdlg | 3.47 | Prairie ecotone |
| *Prunus virginiana* (Rosaceae) | Understory tree | Widespread | A Sap Sdlg | 38.42 | Canyon bottoms |
| *Quercus macrocarpa* (Fagaceae) | Canopy tree | Central | A Sap Sdlg | 69.25 | Upland forest |
| *Rhus aromatica* (Anacardiaceae) | Clonal shrub | Widespread | Sap Sdlg | 0.07 | Canyon bottoms |
| *Rhus glabra* (Anacardiaceae) | Clonal shrub | Widespread | Sap Sdlg | 16.30 | Upper canyon |
| *Ribes americanum* (Grossulariaceae) | Shrub | Boreal | Sdlg | — | Canyon bottoms |
| *Rosa arkansana* (Rosaceae) | Shrub | Central | Sdlg | — | Prairie ecotone |
| *Rubus occidentalis* (Rosaceae) | Shrub | Eastern | Sdlg | — | Upper canyon |
| *Salix amygdaloides* (Salicaceae) | Understory tree | Widespread | Sap Sdlg | 3.39 | Floodplain |
| *Salix eriocephala* (Salicaceae) | Understory tree | Eastern | Sap Sdlg | 0.88 | Floodplain |
| *Salix interior* (Salicaceae) | Understory tree | Central | Sap Sdlg | 4.28 | Floodplain |
| *Smilax lasioneura* (Smilaceae) | Liana | Widespread | Sdlg | — | Upland forest |
| *Symphoricarpos occidentalis* (Caprifoliaceae) | Shrub | Widespread | Sdlg | — | Floodplain |

| | | | | | |
|---|---|---|---|---|---|
| *Tilia americana* (Malvaceae) | Canopy tree | Eastern | A Sap Sdlg | 34.14 | Upland forest |
| *Toxicodendron radicans* (Anacardiaceae) | Shrub/liana | Eastern | Sdlg | — | Upper canyon |
| *Ulmus americana* (Ulmaceae) | Canopy tree | Eastern | A Sap | 1.84 | Upland forest |
| *Ulmus laevis* (Ulmaceae) | Canopy tree | Non-native | Sap | 0.07 | Upper canyon |
| *Ulmus rubra* (Ulmaceae) | Canopy tree | Eastern | A Sap Sdlg | 10.32 | Upland forest |
| *Vitis riparia* (Vitaceae) | Liana | Eastern | A Sap Sdlg | 17.40 | Canyon bottoms & Upland forest |
| *Zanthoxylum americanum* (Rutaceae) | Understory tree | Eastern | Sdlg | — | Upland forest |

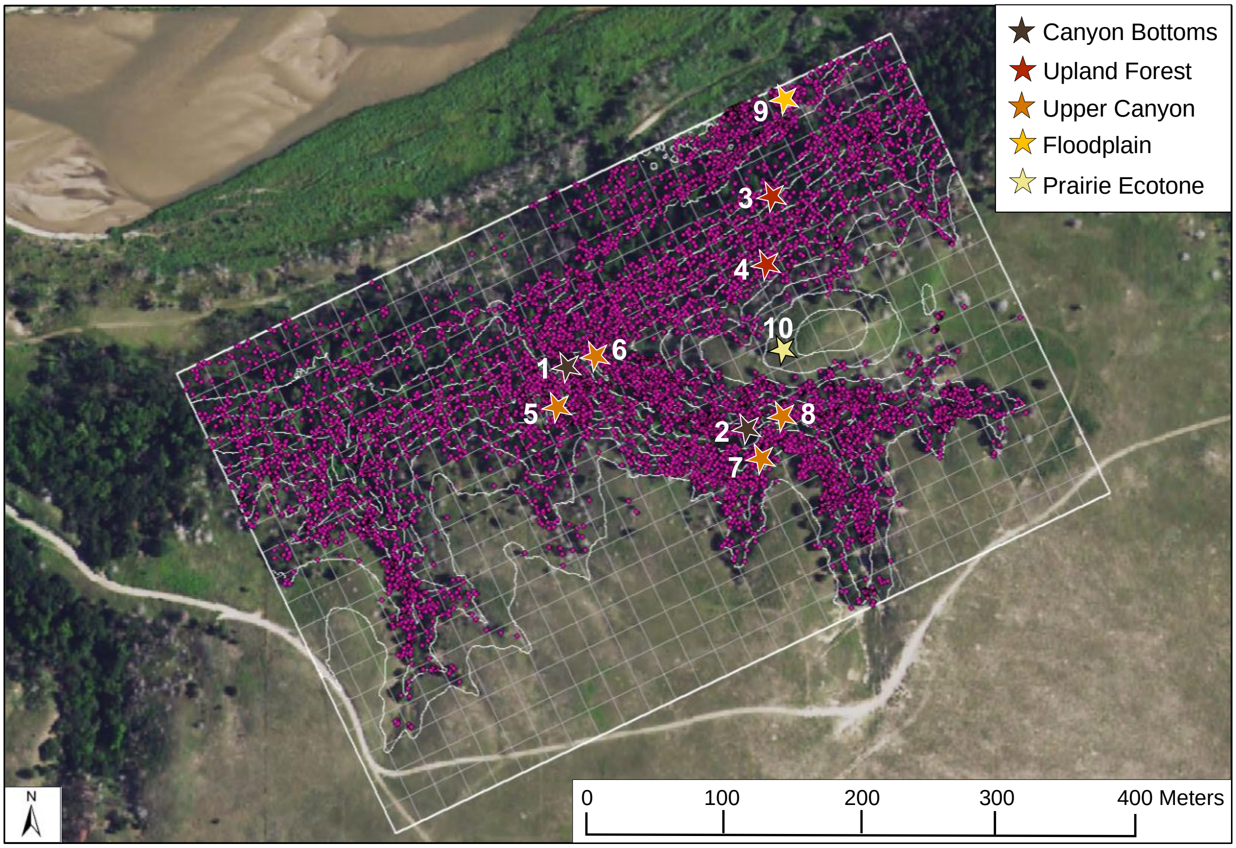


Canyon Bottoms
Lower (1) – 656.2 m, 35.6%, 228°
Upper (2) – 672.4 m, 34.2%, 206°

Upland Forest
Lower (3) – 659.5 m, 26.4%, 319°
Upper (4) – 679.4 m, 28.9%, 324°

Upper Canyon
Lower Northeast (5) – 678.3 m, 27.3%, 266°
Lower Southwest (6) – 660.2 m, 38.6%, 194°
Upper Northeast (7) – 680.6 m, 36.3%, 312°
Upper Southwest (8) – 686.2 m, 30.4%, 174°

Floodplain (9)
– 645.1 m, 7.2%, 234°

Prairie Ecotone (10)
– 696.8 m, 14.9%, 249°

**Figure S1. Locations of the ten microclimate monitoring stations in the Niobrara plot**. A contour plot (5-m elevation change between contours) overlays the Niobrara plot (560 m x 360 m, rectangular boundary indicated by thicker white line), gridlines (finer white lines) indicate the 20 m x 20 m quadrats, and each pink dot represents a mapped woody individual (DBH ≥ 1 cm), all overlain on an aerial image. Numbered stars indicate the locations of microclimate stations and are color-coded by the habitat (Figure 1f) in which they are located. The mean quadrat elevation (m), topographic slope (%), and aspect (°) for each monitoring station is indicated below the map. Sampling efforts were more intensive along the upper canyon slopes to capture the variation in conditions on the Northeast-facing (Stations 5 and 7) versus Southwest-facing (Stations 6 and 8) slopes, and from the high to low end of the main canyon (see map of cumulative annual solar radiation and thermal image of plot; Figure 1d,e). Stations 1 and 2 were near the stream at the canyon bottom, and Station 6 was near a spring. The aerial image is a high-resolution RGBN image obtained on 18 September 2019 with an UltraCam Eagle camera (Vexcel Imaging).

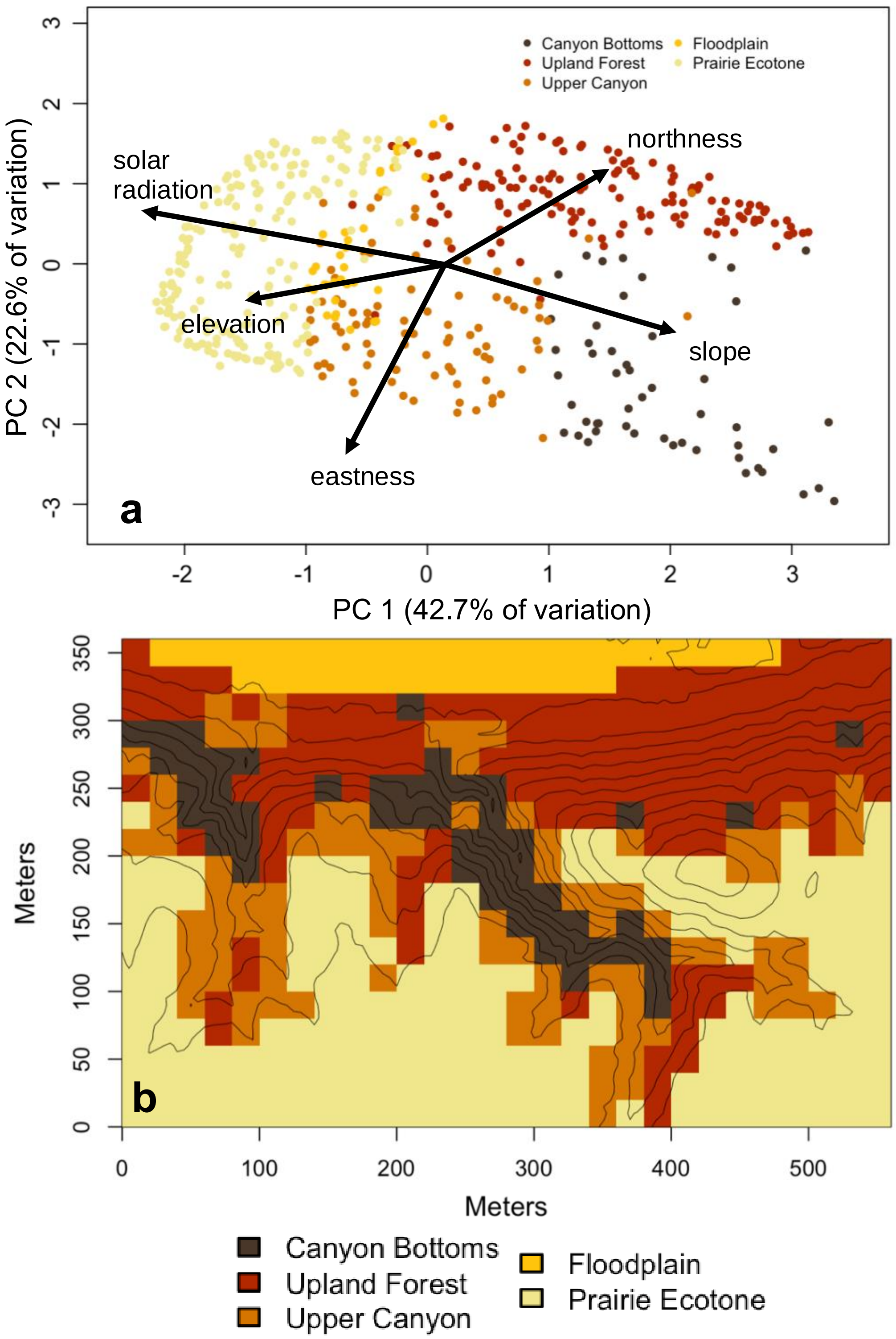


**Figure S2. Derivation of categorical habitats based on topographic variables obtained from a digital elevation model (DEM) of the Niobrara plot.** (a) Biplot of the first two principal components with points (20 x 20 m quadrats) colored by habitat, with arrows depicting the loadings of the topographic variables onto the PC axes; (b) a map of habitats in the Niobrara plot, overlaid with 5-m elevation contour lines. See Table S2 for the loadings of each variable onto PC1 and PC2, and Appendix S1 and Tables S5 for additional details on habitat classification method and habitat types

**Table S2. Loadings of five topographic variables onto the first five principal components (PCs) of a principal components analysis used to define five categorical habitats across the Niobrara plot.** Aspect is decomposed into northness and eastness. Topographic variables were derived from a digital elevation model. Cutoffs of PC1 and PC2 were used to define the habitats, and values less and greater than |0.4| are shown in bold. See *Methods* and Figure S2 for details.

| **Topographic variable** | **PC1** | **PC2** | **PC3** | **PC4** | **PC5** |
|---|---|---|---|---|---|
| **Solar radiation (Wh/m$^2$)** | **-0.61732** | 0.24194 | -0.04688 | 0.06493 | **-0.74430** |
| **Elevation (m)** | -0.37240 | -0.12680 | **0.77426** | **-0.46247** | 0.17855 |
| **Slope (%)** | **0.56816** | -0.29222 | 0.09380 | **-0.45663** | **-0.61196** |
| **Eastness** | -0.14922 | **-0.82778** | 0.12518 | **0.51490** | -0.10828 |
| **Northness** | 0.36765 | 0.39342 | **0.61144** | **0.55523** | -0.16712 |

**Table S3. Summary of habitat-related variation in microclimate in the Niobrara plot.** Microclimate data for each habitat were collected at ten monitoring stations from April to November 2021: Canyon Bottoms, Stations 1 and 2; Upland Forest, Stations 3 and 4; Upper Canyon, Stations 5-8; Floodplain, Station 9; and Prairie Ecotone, Station 10 (Figure S1). Values are the within-habitat average of the within-station average of the daily means across all measurement days of the growing season, except for photosynthetic photon flux density (PPFD), which is the average of the daily total PPFD across all measurement days. All values are shown ± standard deviation, and the average daily minimums and maximums for air temperature (air temp), relative humidity (RH), soil temperature (soil temp), and volumetric water content (VWC) are in parentheses in the second row. Vapor pressure deficit (VPD) was calculated from air temp and RH. Air temp and RH sensors were not deployed until 21 May 2021 for Stations 3, 7, and 9, and until 12 June 2021 for Stations 1, 4, 5, 6, and 8.

| **Habitat** | **PPFD ($mmol/m^2/d$)** | **Air Temp (ºC)** | **RH (%)** | **VPD (kPa)** | **Soil Temp (ºC)** | **VWC (%)** |
|---|---|---|---|---|---|---|
| **Canyon Bottoms** | 842.1 ± 325.2 | 14.8 ± 2.9<br>(-6.9-36.8) | 82.7 ± 8.3<br>(20.0-100) | 0.30 ± 0.2 | 14.4 ± 1.5<br>(3.0-21.7) | 30.3 ± 0.9<br>(26.9-34.3) |
| **Upland Forest** | 370.6 ± 124.8 | 16.9 ± 3.0<br>(-7.1-38.7) | 71.8 ± 10.4<br>(0.0-100) | 0.61 ± 0.3 | 15.6 ± 1.9<br>(-0.03-25.3) | 22.3 ± 0.7<br>(19.6-26.6) |
| **Upper Canyon** | 644.9 ± 203.1 | 17.0 ± 3.3<br>(-7.7-42.9) | 72.0 ± 10.7<br>(13.7-100) | 0.62 ± 0.3 | 15.9 ± 1.7<br>(2.8-26.0) | 23.3 ± 0.4<br>(17.7-37.5) |
| **Flood-plain** | 4337.7 ± 1181.8 | 16.4 ± 2.9<br>(-8.4-39.3) | 83.1 ± 6.2<br>(17.7-100) | 0.36 ± 0.2 | 15.8 ± 1.7<br>(3.2-23.6) | 26.7 ± 0.6<br>(20.7-34.8) |
| **Prairie Ecotone** | 5270.2 ± 1474.3 | 16.4 ± 3.9<br>(-7.4-41.0) | 66.4 ± 11.6<br>(15.9-100) | 0.74 ± 0.4 | 20.9 ± 2.9<br>(2.1-37.1) | 20.1 ± 0.6<br>(18.2-22.2) |

**Table S4. List of 59 numbered candidate generalized additive models (GAMs) for predicting tree (diameter at breast height (DBH) ≥ 1 cm) stem density, basal area, aboveground biomass, and species richness, and seedling (DBH < 1 cm) stem density and species richness in the Niobrara plot.** The two-dimensional splines on x and y, "s(x,y, bs = "ts")", are smoothing terms that penalize the null space and allow for the term to shrink to zero if necessary. Std.elev = standardized elevation (m); northness = cosine(aspect in radians); eastness = sin(aspect in radians); solarMean = cumulative annual solar radiation ($Wh/m^2$); slopeMean = slope (%); "*" represents interactions between covariates, and models including more than one two-way interaction are indicated by covariates in parentheses and squared.

| Number | Model Form |
|---|---|
| 1 | ~ 1 |
| 2 | ~ 1 + s(x,y,bs="ts") |
| 3 | ~ std.elev + s(x,y,bs="ts") |
| 4 | ~ northness + s(x,y,bs="ts") |
| 5 | ~ eastness + s(x,y,bs="ts") |
| 6 | ~ solarMean + s(x,y,bs="ts") |
| 7 | ~ slopeMean + s(x,y,bs="ts") |
| 8 | ~ std.elev + northness + s(x,y,bs="ts") |
| 9 | ~ std.elev + eastness + s(x,y,bs="ts") |
| 10 | ~ std.elev + solarMean + s(x,y,bs="ts") |
| 11 | ~ std.elev + slopeMean + s(x,y,bs="ts") |
| 12 | ~ northness + eastness + s(x,y,bs="ts") |
| 13 | ~ northness + solarMean + s(x,y,bs="ts") |
| 14 | ~ northness + slopeMean + s(x,y,bs="ts") |
| 15 | ~ eastness + solarMean + s(x,y,bs="ts") |
| 16 | ~ eastness + slopeMean + s(x,y,bs="ts") |
| 17 | ~ solarMean + slopeMean + s(x,y,bs="ts") |
| 18 | ~ std.elev + northness + eastness + s(x,y,bs="ts") |
| 19 | ~ std.elev + northness + solarMean + s(x,y,bs="ts") |

| | |
|---|---|
| 20 | ~ std.elev + northness + slopeMean + s(x,y,bs="ts") |
| 21 | ~ std.elev + eastness + solarMean + s(x,y,bs="ts") |
| 22 | ~ std.elev + eastness + slopeMean + s(x,y,bs="ts") |
| 23 | ~ std.elev + solarMean + slopeMean + s(x,y,bs="ts") |
| 24 | ~ northness + eastness + solarMean + s(x,y,bs="ts") |
| 25 | ~ northness + eastness + slopeMean + s(x,y,bs="ts") |
| 26 | ~ northness + solarMean + slopeMean + s(x,y,bs="ts") |
| 27 | ~ eastness + solarMean + slopeMean + s(x,y,bs="ts") |
| 28 | ~ std.elev + northness + eastness + solarMean + s(x,y,bs="ts") |
| 29 | ~ std.elev + northness + eastness + slopeMean + s(x,y,bs="ts") |
| 30 | ~ std.elev + northness + solarMean + slopeMean + s(x,y,bs="ts") |
| 31 | ~ std.elev + eastness + solarMean + slopeMean + s(x,y,bs="ts") |
| 32 | ~ northness + eastness + solarMean + slopeMean + s(x,y,bs="ts") |
| 33 | ~ std.elev + northness + eastness + solarMean + slopeMean + s(x,y,bs="ts") |
| 34 | ~ std.elev * northness + s(x,y,bs="ts") |
| 35 | ~ std.elev * eastness + s(x,y,bs="ts") |
| 36 | ~ std.elev * solarMean + s(x,y,bs="ts") |
| 37 | ~ std.elev * slopeMean + s(x,y,bs="ts") |
| 38 | ~ northness * eastness + s(x,y,bs="ts") |
| 39 | ~ northness * solarMean + s(x,y,bs="ts") |
| 40 | ~ northness * slopeMean + s(x,y,bs="ts") |
| 41 | ~ eastness * solarMean + s(x,y,bs="ts") |
| 42 | ~ eastness * slopeMean + s(x,y,bs="ts") |
| 43 | ~ solarMean * slopeMean + s(x,y,bs="ts") |
| 44 | ~ (std.elev + northness + eastness)^2 + s(x,y,bs="ts") |
| 45 | ~ (std.elev + northness + solarMean)^2 + s(x,y,bs="ts") |
| 46 | ~ (std.elev + northness + slopeMean)^2 + s(x,y,bs="ts") |
| 47 | ~ (std.elev + eastness + solarMean)^2 + s(x,y,bs="ts") |
| 48 | ~ (std.elev + eastness + slopeMean)^2 + s(x,y,bs="ts") |
| 49 | ~ (std.elev + solarMean + slopeMean)^2 + s(x,y,bs="ts") |
| 50 | ~ (northness + eastness + solarMean)^2 + s(x,y,bs="ts") |

| | |
|---|---|
| 51 | ~ (northness + eastness + slopeMean)^2 + s(x,y,bs="ts") |
| 52 | ~ (northness + solarMean + slopeMean)^2 + s(x,y,bs="ts") |
| 53 | ~ (eastness + solarMean + slopeMean)^2 + s(x,y,bs="ts") |
| 54 | ~ (std.elev + northness + eastness + solarMean)^2 + s(x,y, bs="ts") |
| 55 | ~ (std.elev + northness + eastness + slopeMean)^2 + s(x,y, bs="ts") |
| 56 | ~ (std.elev + northness + solarMean + slopeMean)^2 + s(x,y, bs="ts") |
| 57 | ~ (std.elev + eastness + solarMean + slopeMean)^2 + s(x,y, bs="ts") |
| 58 | ~ (northness + eastness + solarMean + slopeMean)^2 + s(x,y, bs="ts") |
| 59 | ~ (std.elev + northness + eastness + solarMean + slopeMean)^2 + s(x,y, bs="ts") |

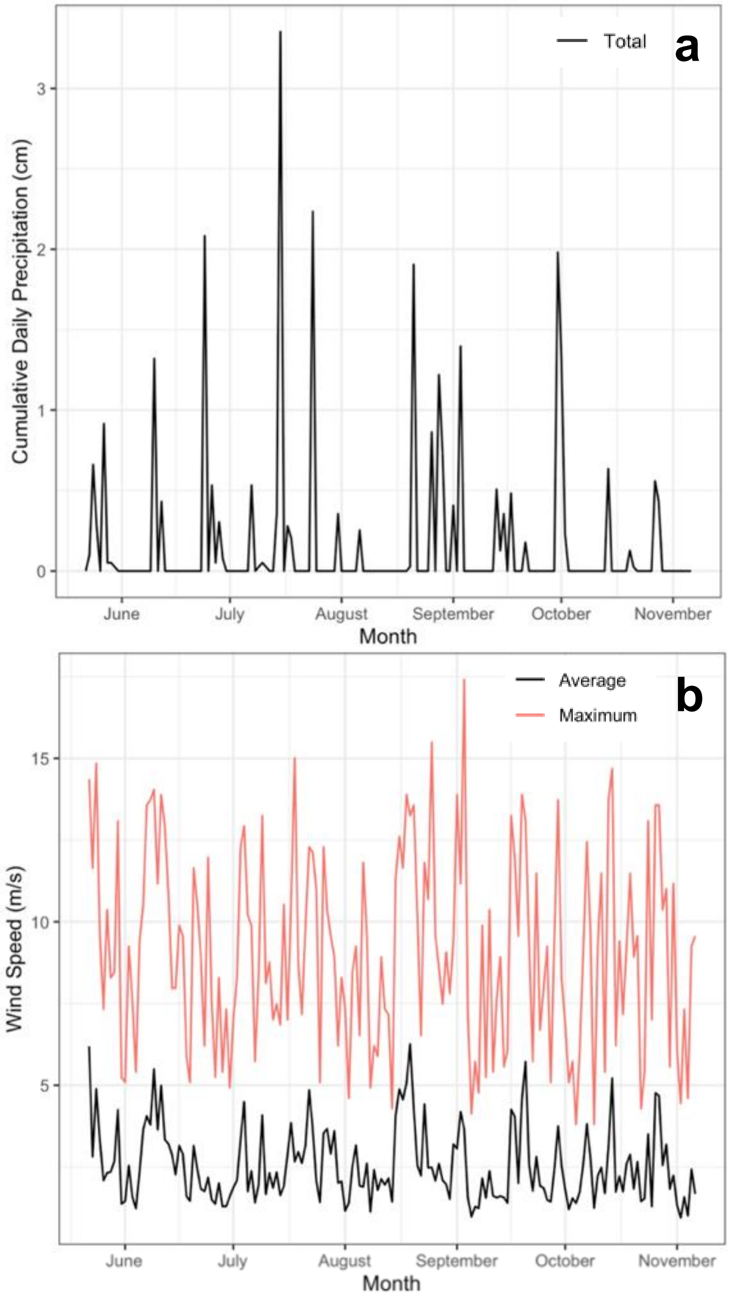


**Figure S3. Cumulative precipitation and variation in wind speed in the prairie ecotone habitat in the Niobrara plot. (**a) Cumulative daily precipitation (cm), and (b) average (black line) and maximum (red line) daily wind speed (m/s) between 21 May-6 November 2021.

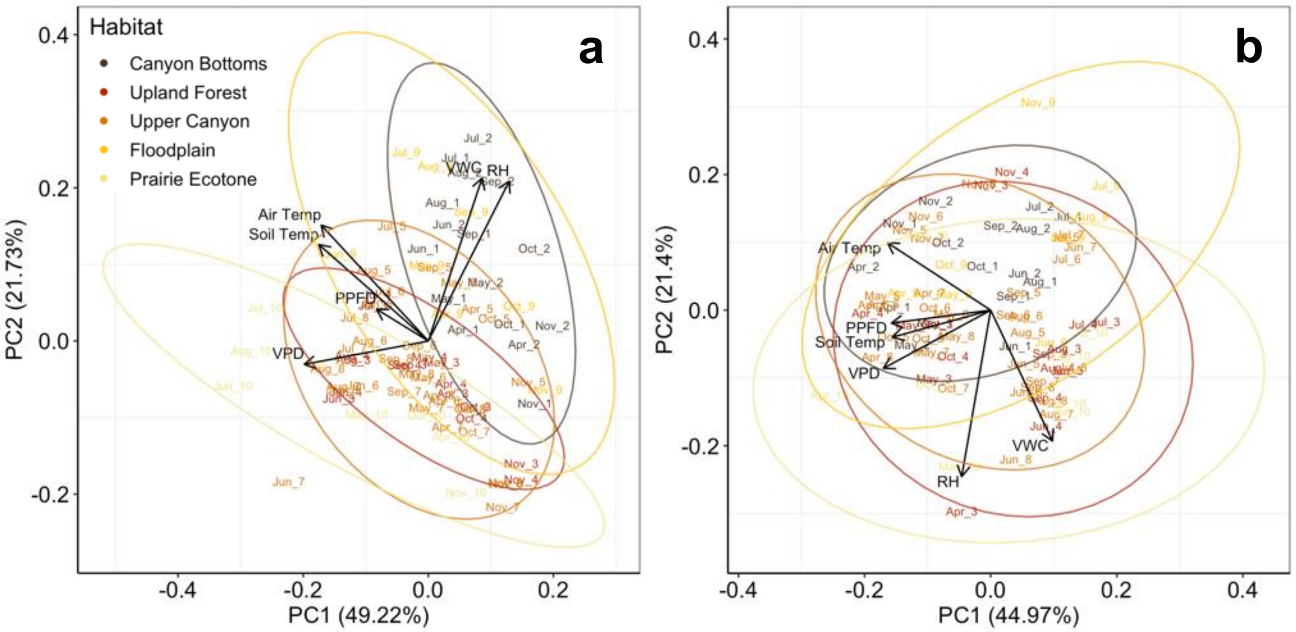


**Figure S4. Multivariate habitat-related microclimate variation in the Niobrara plot, based on data collected at ten monitoring stations from April-November 2021.** Principal Component 1 versus 2 (percent variance explained) from separate Principal Components Analyses on the monthly (a) means and (b) coefficients of variation (CVs) of the microclimatic variables shown in Figure 2. Each point represents a Month-Station (numbers are station numbers), colors indicate habitats, and ellipses are the 95% confidence intervals based on the standard error for each habitat. Means and CVs were imputed for the missing air temperature, relative humidity, and vapor pressure deficit values in April and May, as sensors measuring these variables were not deployed until 21 May 2021 for Stations 3, 7, and 9, and until 12 June 2021 for Stations 1, 4, 5, 6, and 8. Both the effect of habitat type and the interaction between habitat type and month were significant in the perMANOVA on the microclimate means (habitat: $F_{4,40} = 52.7$, $R^2 = 0.39$, $p = 0.001$; habitat × month: $F_{35,40} = 8.4$, $R^2 = 0.54$, $p = 0.001$) and the microclimate CVs (habitat: $F_{4,40} = 11.2$, $R^2 = 0.11$, $p = 0.001$; habitat × month: $F_{35,40} = 9.4$, $R^2 = 0.80$, $p = 0.001$). PPFD = photosynthetic photon flux density (mmol/m$^2$/day); air temperature = air temperature (°C) (c) daily mean); RH = relative humidity (%); VPD = vapor pressure deficit (kPa); soil temp = soil temperature (°C); VWC = volumetric water content (%).

**Table S5. Summary of forest structure, diversity, and topographic variation in the five habitats in the Niobrara plot.** Metrics of forest structure for adults and saplings are on a per hectare basis, and diversity is per quadrat (400 $m^2$), whereas these metrics are per subplot (1 $m^2$) for seedlings. Numbers reported are mean ± standard error for metrics of structure and diversity and are quadrat-level mean (range) for metrics of environmental variation. See Appendix S1 and Figure S2 for details about habitat classification.

| **Variable** | **Size class** | **Canyon bottoms** | **Upland forest** | **Upper canyon** | **Floodplain** | **Prairie ecotone** |
|---|---|---|---|---|---|---|
| **Stem density** | Adult | 561.8 ± 41.7 | 385.8 ± 22.7 | 375.7 ± 39.4 | 185.6 ± 24.4 | 139.2 ± 40.7 |
| | Sapling | 532.0 ± 62.4 | 228.6 ± 24.6 | 413.1 ± 57.8 | 566.2 ± 256.1 | 143.7 ± 33.3 |
| | Seedling | 12.1 ± 1.7 | 13.3 ± 1.4 | 8.3 ± 1.5 | 12.2 ± 1.8 | 6.0 ± 0.7 |
| **Basal area ($m^2$)** | Adult | 25.5 ± 1.77 | 24.7 ± 1.20 | 16.8 ± 1.43 | 14.1 ± 1.91 | 6.7 ± 1.34 |
| | Sapling | 0.39 ± 0.04 | 0.19 ± 0.02 | 0.29 ± 0.04 | 0.32 ± 0.08 | 0.14 ± 0.04 |
| | Seedling | — | — | — | — | — |
| **AGB (Mg)** | Adult | 163.33 ± 12.32 | 171.27 ± 8.84 | 110.548 ± 9.54 | 112.20 ± 15.68 | 42.22 ± 8.76 |
| | Sapling | 0.78 ± 0.09 | 0.40 ± 0.50 | 0.55 ± 0.07 | 1.22 ± 0.33 | 0.33 ± 0.09 |
| | Seedling | — | — | — | — | — |
| **Species richness** | Adult | 5.2 ± 0.2 | 4.0 ± 0.1 | 2.9 ± 0.2 | 2.2 ± 0.2 | 1.5 ± 0.1 |
| | Sapling | 4.5 ± 0.4 | 2.2 ± 0.1 | 2.6 ± 0.3 | 2.2 ± 0.5 | 1.2 ± 0.1 |
| | Seedling | 3.5 ± 0.4 | 1.4 ± 0.2 | 2.2 ± 0.3 | 2.8 ± 0.3 | 1.3 ± 0.1 |
| **Solar radiation ($Wh/m^2$)** | — | 847759 (598604-1079966) | 1006977 (812452-1206939) | 1091673 (836318-1353250) | 1188999 (1139485-1209570) | 1228649 (1112598-1379424) |
| **Elevation (m)** | — | 670.8 | 666.5 | 680.6 | 646.7 | 690.0 |

| | | | | | | |
|---|---|---|---|---|---|---|
| | | (649.8-687.7) | (646.0-693.3) | (651.0-695.5) | (644.8-649.7) | (682.1-698.5) |
| **Slope (%)** | — | 32.7 | 20.7 | 21.3 | 4.7 | 13.5 |
| | | (22.5-41.0) | (6.1-32.5) | (9.3-38.3) | (1.7-7.2) | (1.9-27.9) |
| **Eastness** | — | 0.17 | -0.83 | 0.11 | -0.18 | -0.24 |
| | | (-0.96-1.00) | (-1.00-0.24) | (-0.97-1.00) | (-1.00-0.99) | (-1.0-0.90) |
| **Northness** | — | -0.57 | 0.19 | -0.57 | -0.72 | -0.66 |
| | | (-1.00-0.67) | (-0.97-0.91) | (-1.00-0.74) | (-1.00-0.04) | (-1.00-0.38) |
| **PC1** | — | 1.93 | 1.37 | -0.017 | -0.55 | -1.42 |
| | | (1.02-3.35) | (-0.62-3.14) | (-0.96-2.18) | (-0.98-0.13) | (-2.23- -0.20) |
| **PC2** | — | -1.53 | 0.84 | -0.76 | 0.28 | 0.10 |
| | | (-2.96-0.17) | (-0.64-1.72) | (-2.17-0.88) | (-0.82-1.81) | (-1.35-1.63) |

**Table S6.** ***F*****-values, degrees of freedom (DF),** $\boldsymbol{R^2}$ **values, and** ***p*****-values for the linear models testing variation in tree (diameter at breast height (DBH) ≥ 1 cm) stem density, basal area, aboveground biomass (AGB), species richness, and Shannon's diversity index and seedling (DBH <1 cm) stem density, species richness, and Shannon's diversity index among habitat types in the Niobrara plot**. Significant effects at the $\alpha$ = 0.05 level shown in bold.

| **Response Variable** | ***F*-value** | **DF (Error, Total)** | $\boldsymbol{R^2}$ | ***p*-value** |
|---|---|---|---|---|
| Tree stem density | 37.3 | 4, 334 | 0.30 | **<0.001** |
| Tree basal area | 31.9 | 4, 334 | 0.27 | **<0.001** |
| Tree AGB | 28.9 | 4, 334 | 0.25 | **<0.001** |
| Tree species richness | 58.7 | 4, 334 | 0.41 | **<0.001** |
| Tree Shannon's diversity | 55.4 | 4, 334 | 0.39 | **<0.001** |
| Seedling stem density | 8.4 | 4, 285 | 0.09 | **<0.001** |
| Seedling species richness | 28.4 | 4, 285 | 0.27 | **<0.001** |
| Seedling Shannon's diversity | 26.9 | 4, 285 | 0.26 | **<0.001** |

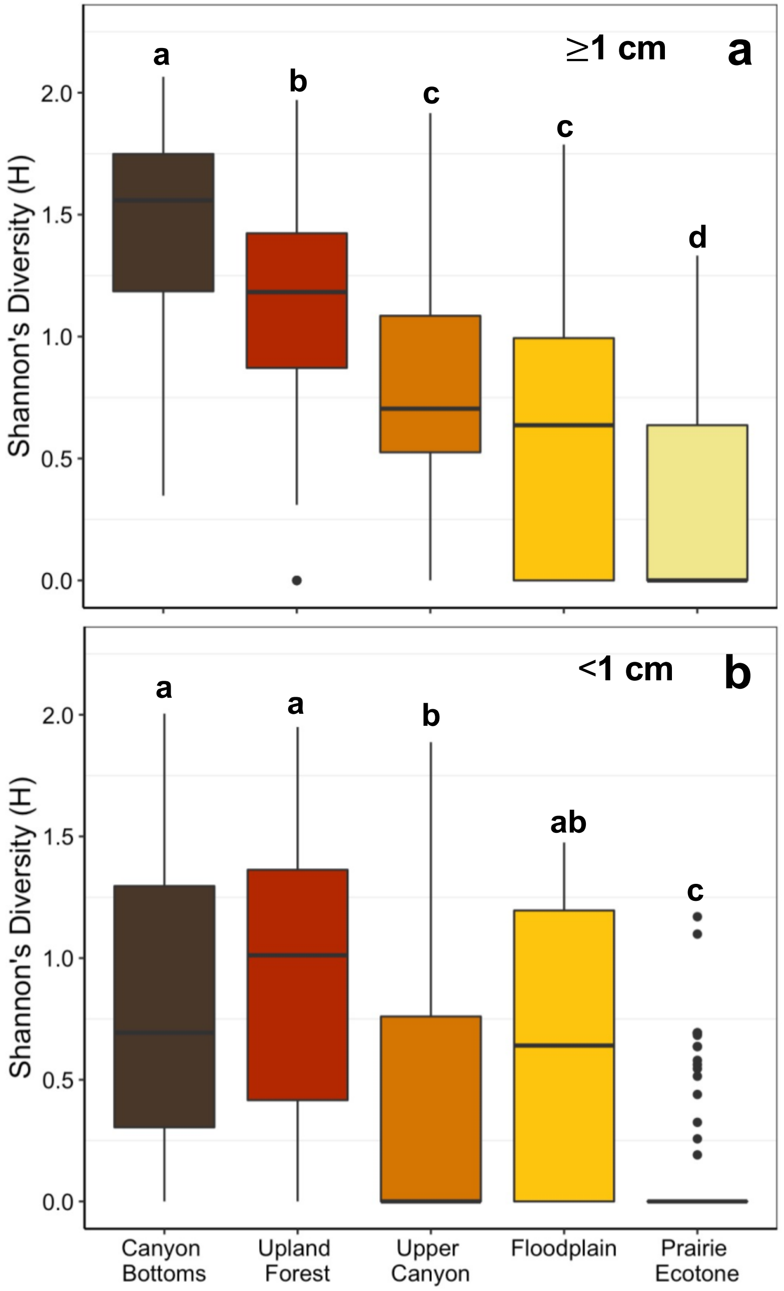


**Figure S5. Variation in Shannon's diversity indices (H) among five habitats in the Niobrara plot.** Boxplots show habitat-related variation for (a) tree (diameter at breast height (DBH) ≥1 cm) diversity and (b) seedling (DBH <1 cm) diversity. Habitats are ordered by increasing light intensity and exposure from canyon bottoms to prairie ecotone (Figure 1f). Letters above the boxes indicate significant pairwise differences at the $\alpha$ = 0.05 level after correction for multiple comparisons using Tukey's Honestly Significant Differences. Boxes indicate the median (center line) and first and third quartiles; lower and upper whiskers indicate the first and third quartiles ±1.5 times the interquartile range; points indicate outliers.

**Table S7. Abundance of the 37 woody species (27 reaching diameter at breast height (DBH) ≥1 cm) in five habitats in the Niobrara plot**. Habitats are ordered by increasing exposure, the habitat with the highest combined abundance of adults, saplings, and seedlings per species is shown in bold, and the DBH (cm) range is shown for species reaching DBH ≥1 cm.

| **Species** | **Size class** | **Canyon bottoms** | **Upland forest** | **Upper canyon** | **Flood-plain** | **Prairie ecotone** | **DBH range** |
|---|---|---|---|---|---|---|---|
| *Acer negundo* | Total | **3** | 0 | 0 | 1 | 0 | 1.0-35.6 |
| | Adult | **2** | 0 | 0 | 0 | 0 | 11.9-35.6 |
| | Sapling | **1** | 0 | 0 | 1 | 0 | 1.0-3.0 |
| | Seedling | **0** | 0 | 0 | 0 | 0 | NA |
| *Amorpha canescens* | Total | 0 | **8** | 0 | 0 | 0 | NA |
| | Adult | 0 | **0** | 0 | 0 | 0 | |
| | Sapling | 0 | **0** | 0 | 0 | 0 | |
| | Seedling | 0 | **8** | 0 | 0 | 0 | |
| *Betula papyrifera* | Total | **18** | 8 | 3 | 0 | 0 | 1.5-39.4 |
| | Adult | **15** | 7 | 3 | 0 | 0 | 5.4-39.4 |
| | Sapling | **3** | 0 | 0 | 0 | 0 | 1.5-5.0 |
| | Seedling | **0** | 1 | 0 | 0 | 0 | NA |
| *Celastrus scandens* | Total | **19** | 14 | 17 | 0 | 0 | 1.0-1.5 |
| | Adult | **0** | 0 | 0 | 0 | 0 | NA |
| | Sapling | **6** | 2 | 0 | 0 | 0 | 1.0-1.5 |
| | Seedling | **13** | 12 | 17 | 0 | 0 | NA |
| *Celtis occidentalis* | Total | 170 | **379** | 90 | 29 | 2 | 1.0-61.0 |
| | Adult | 36 | **85** | 17 | 5 | 0 | 5.2-61.0 |
| | Sapling | 69 | **41** | 19 | 5 | 0 | 1.0-4.9 |
| | Seedling | 65 | **253** | 54 | 19 | 2 | NA |
| *Cornus sericea* | Total | **2** | 0 | 0 | 0 | 0 | 1.0-1.0 |
| | Adult | **0** | 0 | 0 | 0 | 0 | NA |
| | Sapling | **2** | 0 | 0 | 0 | 0 | 1.0-1.0 |

| | | | | | | | |
|---|---|---|---|---|---|---|---|
| | Seedling | **0** | 0 | 0 | 0 | 0 | NA |
| *Dalea villosa* | Total | 0 | 0 | 0 | 0 | **2** | NA |
| | Adult | 0 | 0 | 0 | 0 | **0** | |
| | Sapling | 0 | 0 | 0 | 0 | **0** | |
| | Seedling | 0 | 0 | 0 | 0 | **2** | |
| *Fraxinus pennsylvanica* | Total | 76 | **220** | 55 | 200 | 3 | 1.0-80.1 |
| | Adult | 37 | **81** | 27 | 110 | 1 | 5.1-80.1 |
| | Sapling | 30 | **19** | 14 | 89 | 2 | 1.0-4.9 |
| | Seedling | 9 | **120** | 14 | 1 | 0 | NA |
| *Juglans nigra* | Total | 8 | **23** | 1 | 0 | 0 | 2.0-74.0 |
| | Adult | 4 | **16** | 1 | 0 | 0 | 5.9-74.0 |
| | Sapling | 3 | **3** | 0 | 0 | 0 | 2.0-3.8 |
| | Seedling | 1 | **4** | 0 | 0 | 0 | NA |
| *Juniperus virginiana* | Total | 740 | 966 | **1032** | 14 | 189 | 1.0-68.6 |
| | Adult | 458 | 647 | **573** | 5 | 126 | 5.1-68.6 |
| | Sapling | 256 | 254 | **447** | 2 | 62 | 1.0-5.0 |
| | Seedling | 26 | 65 | **12** | 7 | 1 | NA |
| *Lonicera dioica* | Total | **14** | 0 | 0 | 0 | 0 | NA |
| | Adult | **0** | 0 | 0 | 0 | 0 | |
| | Sapling | **0** | 0 | 0 | 0 | 0 | |
| | Seedling | **14** | 0 | 0 | 0 | 0 | |
| *Morus alba* | Total | **1** | 0 | 0 | 0 | 0 | 14.0* |
| | Adult | **1** | 0 | 0 | 0 | 0 | 14.0 |
| | Sapling | **0** | 0 | 0 | 0 | 0 | NA |
| | Seedling | **0** | 0 | 0 | 0 | 0 | NA |
| *Ostrya virginiana* | Total | 640 | **938** | 203 | 9 | 0 | 1.0-17.6 |
| | Adult | 287 | **439** | 90 | 4 | 0 | 5.1-17.6 |
| | Sapling | 289 | **354** | 102 | 4 | 0 | 1.0-5.0 |
| | Seedling | 64 | **145** | 11 | 1 | 0 | NA |
| *Parthenocissus quinquefolia* | Total | 17 | **47** | 12 | 11 | 0 | 1.0-3.4 |
| | Adult | 0 | **0** | 0 | 0 | 0 | NA |

| | | | | | | | |
|---|---|---|---|---|---|---|---|
| | Sapling | 5 | **6** | 4 | 9 | 0 | 1.0-3.4 |
| | Seedling | 12 (23%)† | **41 (30%)**† | 8 (10%)† | 2 (6%)† | 0 | NA |
| *Physocarpus opulifolius* | Total | **4** | 1 | 0 | 3 | 0 | 1.3-1.6 |
| | Adult | **0** | 0 | 0 | 0 | 0 | NA |
| | Sapling | **1** | 1 | 0 | 0 | 0 | 1.3-1.6 |
| | Seedling | **3** | 0 | 0 | 3 | 0 | NA |
| *Pinus ponderosa* | Total | 46 | 63 | **87** | 0 | 25 | 1.0-67.9 |
| | Adult | 41 | 55 | **78** | 0 | 24 | 5.3-67.9 |
| | Sapling | 3 | 8 | **9** | 0 | 1 | 1.0-5.0 |
| | Seedling | 2 | 0 | **0** | 0 | 0 | NA |
| *Populus deltoides* | Total | 1 | 3 | 0 | **185** | 0 | 1.0-85.5 |
| | Adult | 1 | 3 | 0 | **2** | 0 | 5.2-85.5 |
| | Sapling | 0 | 0 | 0 | **182** | 0 | 1.0-3.8 |
| | Seedling | 0 | 0 | 0 | **1** | 0 | NA |
| *Prunus americana* | Total | 0 | 1 | 9 | 25 | **40** | 1.0-4.8 |
| | Adult | 0 | 0 | 0 | 0 | **0** | NA |
| | Sapling | 0 | 1 | 8 | 12 | **26** | 1.0-4.8 |
| | Seedling | 0 | 0 | 1 | 13 | **14** | NA |
| *Prunus virginiana* | Total | **225** | 113 | 224 | 0 | 0 | 1.0-5.7 |
| | Adult | **2** | 0 | 0 | 0 | 0 | 5.7-5.7 |
| | Sapling | **216** | 106 | 197 | 0 | 0 | 1.0-5.0 |
| | Seedling | **7** | 7 | 27 | 0 | 0 | NA |
| *Quercus macrocarpa* | Total | 148 | **408** | 310 | 47 | 66 | 1.1-67.3 |
| | Adult | 129 | **370** | 283 | 37 | 54 | 5.1-67.3 |
| | Sapling | 13 | **16** | 22 | 7 | 8 | 1.1-4.9 |
| | Seedling | 6 | **22** | 5 | 3 | 4 | NA |
| *Rhus aromatica* | Total | 1 | 0 | 0 | 0 | **5** | 1.1* |
| | Adult | 0 | 0 | 0 | 0 | **0** | NA |
| | Sapling | 1 | 0 | 0 | 0 | **0** | 1.1* |
| | Seedling | 0 | 0 | 0 | 0 | **5** | NA |

| | | | | | | | |
|---|---|---|---|---|---|---|---|
| *Rhus glabra* | Total | 34 | 80 | 219 | 3 | **374** | 1.0-3.9 |
| | Adult | 0 | 0 | 0 | 0 | **0** | NA |
| | Sapling | 32 | 38 | 110 | 2 | **39** | 1.0-3.9 |
| | Seedling | 2 | 42 | 109 | 1 | **335** | NA |
| *Ribes americanum* | Total | **1** | 0 | 0 | 0 | 0 | NA |
| | Adult | **0** | 0 | 0 | 0 | 0 | |
| | Sapling | **0** | 0 | 0 | 0 | 0 | |
| | Seedling | **1** | 0 | 0 | 0 | 0 | |
| *Rosa arkansana* | Total | 0 | 2 | 2 | 0 | **58** | NA |
| | Adult | 0 | 0 | 0 | 0 | **0** | |
| | Sapling | 0 | 0 | 0 | 0 | **0** | |
| | Seedling | 0 | 2 | 2 | 0 | **58** | |
| *Rubus occidentalis* | Total | 65 | 131 | **143** | 6 | 17 | NA |
| | Adult | 0 | 0 | **0** | 0 | 0 | |
| | Sapling | 0 | 0 | **0** | 0 | 0 | |
| | Seedling | 65 | 131 | **143** | 6 | 17 | |
| *Salix amygdaloides* | Total | 0 | 0 | 0 | **53** | 0 | 1.0-2.6 |
| | Adult | 0 | 0 | 0 | **0** | 0 | NA |
| | Sapling | 0 | 0 | 0 | **46** | 0 | 1.0-2.6 |
| | Seedling | 0 | 0 | 0 | **7** | 0 | NA |
| *Salix eriocephala* | Total | 0 | 0 | 0 | **17** | 0 | 1.0-3.1 |
| | Adult | 0 | 0 | 0 | **0** | 0 | NA |
| | Sapling | 0 | 0 | 0 | **12** | 0 | 1.0-3.1 |
| | Seedling | 0 | 0 | 0 | **5** | 0 | NA |
| *Salix interior* | Total | 0 | 0 | 0 | **83** | 0 | 1.0-1.7 |
| | Adult | 0 | 0 | 0 | **0** | 0 | NA |
| | Sapling | 0 | 0 | 0 | **58** | 0 | 1.0-1.7 |
| | Seedling | 0 | 0 | 0 | **25** | 0 | NA |
| *Smilax lasioneura* | Total | 0 | **5** | 2 | 0 | 0 | NA |
| | Adult | 0 | **0** | 0 | 0 | 0 | |
| | Sapling | 0 | **0** | 0 | 0 | 0 | |

| | | | | | | | |
|---|---|---|---|---|---|---|---|
| | Seedling | 0 | **5** | 2 | 0 | 0 | |
| *Symphoricarpos occidentalis* | Total | 2 | 8 | 4 | **111** | 8 | NA |
| | Adult | 0 | 0 | 0 | **0** | 0 | |
| | Sapling | 0 | 0 | 0 | **0** | 0 | |
| | Seedling | 2 | 8 | 4 | **111** | 8 | |
| *Tilia americana* | Total | 238 | **548** | 76 | 20 | 1 | 1.0-92.7 |
| | Adult | 91 | **264** | 35 | 14 | 1 | 5.2-92.7 |
| | Sapling | 30 | **20** | 6 | 2 | 0 | 1.0-4.9 |
| | Seedling | 117 | **264** | 35 | 4 | 0 | NA |
| *Toxicodendron radicans* | Total | 63 | 24 | **80** | 18 | 4 | NA |
| | Adult | 0 | 0 | **0** | 0 | 0 | |
| | Sapling | 0 | 0 | **0** | 0 | 0 | |
| | Seedling | 63 | 24 | **80** | 18 | 4 | |
| *Ulmus americana* | Total | 6 | **11** | 3 | 5 | 0 | 1.2-47.3 |
| | Adult | 6 | **7** | 3 | 4 | 0 | 7.1-47.3 |
| | Sapling | 0 | **4** | 0 | 1 | 0 | 1.2-4.1 |
| | Seedling | 0 | **0** | 0 | 0 | 0 | NA |
| *Ulmus laevis* | Total | 0 | 0 | **1** | 0 | 0 | 1.2* |
| | Adult | 0 | 0 | **0** | 0 | 0 | NA |
| | Sapling | 0 | 0 | **1** | 0 | 0 | 1.2* |
| | Seedling | 0 | 0 | **0** | 0 | 0 | NA |
| *Ulmus rubra* | Total | 62 | **69** | 19 | 7 | 0 | 1-61.5 |
| | Adult | 30 | **57** | 14 | 7 | 0 | 5.1-61.5 |
| | Sapling | 22 | **5** | 5 | 0 | 0 | 1.0-4.6 |
| | Seedling | 10 | **7** | 0 | 0 | 0 | NA |
| *Vitis riparia* | Total | 121 | **225** | 44 | 45 | 1 | 1.0-10.9 |
| | Adult | 6 | **6** | 3 | 5 | 0 | 5.2-10.9 |
| | Sapling | 82 | **82** | 31 | 21 | 0 | 1.0-5.0 |
| | Seedling | 33 | **137** | 10 | 19 | 1 | NA |
| *Zanthoxylum americanum* | Total | 0 | **3** | 0 | 1 | 0 | NA |
| | Adult | 0 | **0** | 0 | 0 | 0 | |

| | | | | | |
|---|---|---|---|---|---|
| Sapling | 0 | **0** | 0 | 0 | 0 |
| Seedling | 0 | **3** | 0 | 1 | 0 |

*Indicates only one individual of that species with a diameter ≥1 cm (i.e., no DBH range).

†For *Parthenocissus quinquefolia*, due to difficulty in identifying individuals, seedlings were marked as present/absent in each subplot; these values represent the total number and percent of quadrats per habitat with this species present.

**Table S8. Results from AIC model selection for the top two models, global model, and null model predicting tree (diameter at breast height (DBH) ≥1 cm) stem density, basal area, aboveground biomass (AGB), and species richness, and seedling (DBH <1 cm) stem density and species richness as a function of variation in topographic variables across the Niobrara plot**. K: number of model parameters; weight = calculated Akaike weight; solar: cumulative annual solar radiation (Wh/m$^2$); s(x,y): the two-dimensional thin-plate spline between x and y. Transformations applied to response variables to address non-normality are indicated in parentheses around each variable. The full list of 59 candidate models for each response variable is provided in Table S4.

| **Response Variable** | **Model** | **AIC** | **k** | **ΔAIC** | **Weight** | **Adjusted *R*$^2$** |
|---|---|---|---|---|---|---|
| log(tree stem density) | **Top**: ~ (elevation + eastness + slope)^2 + s(x,y) | 714.9 | 37 | 0 | 0.756 | 0.69 |
| | **2$^{nd}$**: ~ (elevation + northness + eastness + slope)^2 + s(x,y) | 717.4 | 41 | 2.5 | 0.218 | 0.69 |
| | **Global**: ~ (elevation + northness + eastness + solar + slope)^2 + s(x,y) | 751.2 | 24 | 36.3 | 0 | 0.54 |
| | **Null**: ~ 1 | 1086 | 2 | 371.2 | 0 | 0 |
| square root( tree basal area) | **Top**: ~ (elevation + northness + slope)^2 + s(x,y) | 59.64 | 37 | 0 | 0.483 | 0.56 |
| | **2$^{nd}$**: ~ (elevation + northness + eastness + slope)^2 + s(x,y) | 61.74 | 41 | 2.1 | 0.169 | 0.56 |
| | **Global**: ~ (elevation + northness + eastness + solar + slope)^2 + s(x,y) | 73.86 | 27 | 14.2 | 0 | 0.53 |
| | **Null**: ~ 1 | 307.3 | 2 | 247.6 | 0 | 0 |
| square root( tree AGB) | **Top**: ~ (elevation + northness + slope)^2 + s(x,y) | 3103 | 37 | 0 | 0.227 | 0.52 |
| | **2$^{nd}$**: ~ northness * slope + s(x,y) | 3103 | 34 | 0.05 | 0.221 | 0.52 |
| | **Global**: ~ (elevation + northness + eastness + solar + slope)^2 + s(x,y) | 3114 | 27 | 11.3 | 0.001 | 0.49 |
| | **Null**: ~ 1 | 3324 | 2 | 221.2 | 0 | 0 |

| | | | | | | |
|---|---|---|---|---|---|---|
| cubed root(tree species richness) | **Top**: ~ (elevation + northness + slope)^2 + s(x,y) | -211.8 | 37 | 0 | 0.442 | 0.76 |
| | **2nd**: ~ elevation + slope + s(x,y) | -209.2 | 33 | 2.6 | 0.121 | 0.75 |
| | **Global**: ~ (elevation + northness + eastness + solar + slope)^2 + s(x,y) | -178.9 | 27 | 32.8 | 0 | 0.72 |
| | **Null**: ~ 1 | 238.8 | 2 | 450.5 | 0 | 0 |
| log(seedling stem density) | **Top**: ~ elevation + s(x,y) | 798 | 32 | 0 | 0.082 | 0.30 |
| | **2nd**: ~ solar * slope + s(x,y) | 798.3 | 29 | 0.3 | 0.071 | 0.30 |
| | **Global**: ~ (elevation + northness + eastness + solar + slope)^2 + s(x,y) | 804.8 | 28 | 6.8 | 0.003 | 0.28 |
| | **Null**: ~ 1 | 882.4 | 2 | 84.4 | 0 | 0 |
| inverse square root(seedling species richness) | **Top**: ~ elevation + s(x,y) | -175.7 | 32 | 0 | 0.18 | 0.49 |
| | **2nd**: ~ elevation + northness + s(x,y) | -175.1 | 23 | 0.6 | 0.13 | 0.49 |
| | **Global**: ~ (elevation + northness + eastness + solar + slope)^2 + s(x,y) | -152.8 | 39 | 22.9 | 0 | 0.03 |
| | **Null**: ~ 1 | 2.83 | 2 | 178.6 | 0 | 0 |

**Table S9. Summary statistics for the top candidate models of forest structure and diversity as a function of topography across the Niobrara plot.** Parameter estimates, standard errors, *t*-values, and *p*-values for the top candidate models predicting tree (diameter at breast height (DBH) ≥ 1 cm) stem density (adjusted $R^2$ = 0.69), basal area (adjusted $R^2$ = 0.56), aboveground biomass (AGB; adjusted $R^2$ = 0.52), and species richness (adjusted $R^2$ = 0.76), and seedling (DBH <1 cm) stem density (adjusted $R^2$ = 0.30) and species richness (adjusted $R^2$ = 0.49) as a function of variation in topographic variables. Interactions between variables are indicated by "*"; significant effects at the $\alpha$ = 0.05 level shown in bold. Parameters have not been back-transformed. See Table S4 for the full list of models.

| **Response Variable** | **Model Term** | **Estimate** | **Standard Error** | ***t*-value** | ***p*-value** |
|---|---|---|---|---|---|
| Tree stem density | Intercept | 7.215 | 0.490 | 14.717 | **<0.001** |
| | elevation | -0.108 | 0.015 | -7.091 | **<0.001** |
| | eastness | 0.234 | 0.182 | 1.284 | 0.200 |
| | slope | -0.0005 | 0.015 | -0.035 | 0.972 |
| | elevation*eastness | 0.007 | 0.004 | 1.732 | 0.084 |
| | elevation*slope | 0.003 | 0.0004 | 6.309 | **<0.001** |
| | eastness*slope | -0.019 | 0.007 | -2.909 | **0.003** |
| Tree basal area | Intercept | 3.983 | 0.982 | 4.054 | **<0.001** |
| | elevation | -0.080 | 0.031 | -2.596 | **0.010** |
| | northness | -0.455 | 0.437 | -1.041 | 0.299 |
| | slope | 0.052 | 0.029 | 1.798 | 0.073 |
| | elevation*northness | -0.003 | 0.010 | -0.300 | 0.765 |
| | elevation*slope | 0.002 | 0.0008 | 2.375 | **0.018** |
| | northness*slope | 0.042 | 0.016 | 2.695 | **0.007** |
| Tree AGB | Intercept | 10.190 | 2.752 | 3.703 | **<0.001** |
| | elevation | -0.196 | 0.087 | -2.263 | **0.024** |
| | northness | -1.096 | 1.228 | -0.892 | 0.373 |
| | slope | 0.136 | 0.082 | 1.663 | 0.097 |
| | elevation*northness | -0.006 | 0.027 | -0.232 | 0.816 |

| | | | | | |
|---|---|---|---|---|---|
| | elevation*slope | 0.005 | 0.002 | 1.959 | **0.051** |
| | northness*slope | 0.098 | 0.044 | 2.233 | **0.027** |
| Tree species richness | Intercept | 1.630 | 0.126 | 12.984 | <**0.001** |
| | elevation | -0.019 | 0.004 | -4.872 | <**0.001** |
| | northness | -0.117 | 0.058 | -2.012 | **0.045** |
| | slope | 0.021 | 0.004 | 5.506 | <**0.001** |
| | elevation*northness | 0.004 | 0.001 | 2.966 | **0.003** |
| | elevation*slope | 0.00003 | 0.0001 | 0.247 | 0.805 |
| | northness*slope | -0.0006 | 0.002 | -0.278 | 0.781 |
| Seedling stem density | Intercept | 2.656 | 0.375 | 7.075 | <**0.001** |
| | elevation | -0.030 | 0.013 | -2.362 | **0.019** |
| Seedling species richness | Intercept | 0.364 | 0.068 | 5.326 | <**0.001** |
| | elevation | 0.013 | 0.002 | 5.691 | <**0.001** |
